# Insights into the FOXE3 Transcriptional Network and Disease Mechanisms from the Investigation of a Regulatory Variant Driving Complex Microphthalmia

**DOI:** 10.1101/2025.01.13.632782

**Authors:** Julie Plaisancié, Clémentine Angée, Elisa Erjavec, Isabelle Raymond-Letron, Jean-Yves Douet, Mathilde Goetz, Catherine Vincent-Delorme, Ino D. Karemaker, Marijke Baltissen, Michiel Vermeulen, Leonardo Valdivia, Fabienne Jabot-Hanin, Pierre David, Djihad Hadjadj, Yanad Abou Monsef, Faouzi Lyazrhi, Patrick Calvas, Jean-Michel Rozet, Nicolas Chassaing, Lucas Fares-Taie

## Abstract

*FOXE3* encodes a conserved, lens-specific transcription factor essential for eye development. Biallelic mutations in *FOXE3* lead to a spectrum of ocular anomalies, from cataracts to complex microphthalmia (CM), with clinical severity correlating to genotype. In a CM case with a truncating mutation (p.Cys240*), we identified a regulatory variant (rv, rs745674596 G>A) 3 kb upstream of *FOXE3*. Mouse models harboring either the rv or a frameshift mutation were generated in homozygosity (*Foxe3rv/rv, Foxe3-/-*) and compound heterozygosity (*Foxe3rv/Foxe3-*). Phenotypic analysis revealed progressive severity: Foxe3rv/rv mice exhibited cataracts and anterior segment dysgenesis, Foxe3rv/Foxe3-displayed more severe anomalies, and *Foxe3-/-* mice consistently developed CM. These findings align with human genotype-phenotype relationships. Notably, a direct correlation between protein levels and ocular phenotype was observed, with no association to mRNA levels. In Foxe3-/- mice, CM resulted from early disorganization of the anterior lens epithelium, leading to degeneration and ocular involution. Transcription factor binding assays identified USF2 as a key regulator of *FOXE3* expression, positioning USF2 as a promising candidate in ocular development and disease, enhancing our understanding of the FOXE3-related network. This study underscores the importance of integrated approaches to identify genetic variants and cis-regulatory elements, revealing a novel mechanism for microphthalmia through degeneration and involution.

## INTRODUCTION

Vertebrate eye development relies on the precise regulation of gene networks and coordinated interactions among tissues of diverse embryonic origins, which are crucial for forming and patterning fundamental ocular structures (Casey et al. 2023). The lens, a critical component in this process, requires accurate gene expression for proper development (Casey et al. 2023; Cardozo et al. 2023). Multiple transcription factors and signaling pathways orchestrate lens development, including cell proliferation, differentiation, and patterning(Washington et al. 2009).

*FOXE3*, a single-exon gene encoding a 319-amino-acid forkhead family protein, is crucial for lens development (Plaisancié et al. 2018). It regulates early cell differentiation and maintains lens integrity throughout growth by interacting with PAX6 and SOX2 and modulating signaling pathways such as FGF and Wnt, which are essential for lens cell proliferation and patterning (Smith et al. 2009; Cardozo et al. 2023). Pathogenic variants in *FOXE3* are linked to various eye disorders, ranging from congenital cataracts and absence of the lens (aphakia) to anterior segment dysgenesis (ASD) and severe bilateral complex microphthalmia with lens anomalies. These mutations exhibit both autosomal dominant and recessive inheritance patterns, with recessive mutations often leading to severe conditions such as complex microphthalmia and dominant mutations resulting in milder phenotypes like cataracts and ASD (Plaisancié et al. 2018; Reis et al. 2021).

*FOXE3* expression is regulated by several mechanisms, including transcription factors, signaling pathways and epigenetic modifications involving nearby regulatory elements (Valleix et al. 2006; Zhao et al. 2019). Despite the critical role of these mechanisms, the detailed regulatory architecture of *FOXE3* locus and the impact of variants on these regulatory elements are not fully understood.

In this study, we identified a heterozygous pathogenic *FOXE3* nonsense variant (p.Cys240*) (Plaisancié et al. 2018; Reis et al. 2021; Basharat et al. 2023) in a patient with severe bilateral complex microphthalmia, indicative of autosomal recessive inheritance and suggesting the presence of a second loss-of-function allele. Sanger sequencing revealed a non-coding variant in a conserved region located 3 kb upstream of *FOXE3*. This region was previously implicated in mice exhibiting complex microphthalmia and cataract (Wada et al. 2011). Mouse models harboring the non-coding variant, a truncating mutation, or both, demonstrated a genotype-dependent reduction in FOXE3 protein levels. While single heterozygotes exhibited normal ocular development, homozygosity for the non-coding or null allele, as well as compound heterozygosity, resulted in progressively severe ocular defects. Notably, *Foxe3-/-* mice initially developed normal eyes, but these later regressed to complex microphthalmia, underscoring FOXE3’s critical role in maintaining lens development and stability.

Further investigation revealed that the non-coding variant disrupts USF2 binding, and *Usf2* knockdown experiments confirmed its role in downregulating *Foxe3*. These findings highlight the significance of identifying pathogenic non-coding variants in ocular developmental disorders and provide valuable insights into the regulatory architecture of eye development, with genotype-phenotype correlation in partially resolved cases aiding in the discovery of such non-coding mutations.

## RESULTS

### Targeted and whole-genome sequencing revealed a likely disease-causing non-coding variant upstream of *FOXE3* in a patient

The proband, an 11-year-old girl, presented with left complex microphthalmia with Peters’ anomaly and a right polar cataract. She had no significant medical history and was the first child of healthy, young Caucasian parents with no notable family medical history.

Targeted sequencing using a 119-gene panel identified a heterozygous *FOXE3* c.720C>A (NM_012186.3) pathogenic variant, resulting in a p.Cys240* protein change and a premature stop codon. Sanger sequencing-based familial segregation analysis revealed that the pathogenic variant was inherited from the asymptomatic father (**Figure 1A**).

**Figure 1.**
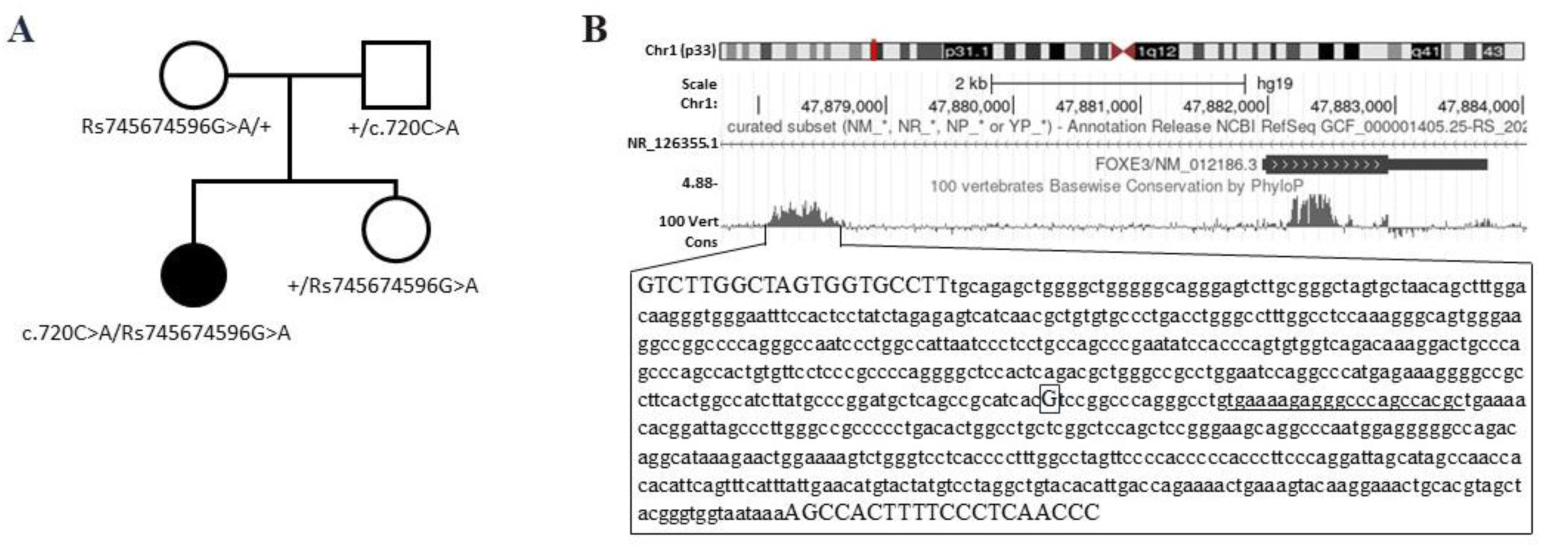
Clinical and Genetic Features of the Proband, Including the Position of the Regulatory Variant in the Highly Conserved Non-Coding Region Upstream of *FOXE3*. **(A)** Family pedigree illustrating the segregation of the c.720C>A nonsense (p.Cys240*) and rs745674596G>A variants. **(B)** UCSC screenshot showing the highly conserved region approximately 3 kb upstream of the *FOXE3* coding sequence, containing the rs745674596G>A variant (highlighted in a square and capitalized). The forward and reverse primers used for PCR amplification are also capitalized. The sequence orthologous to the naturally occurring deletion in the mouse that causes microphthalmia is underlined.

The *FOXE3* c.720C>A nonsense mutation has been reported multiple times, typically with a second mutation in trans in individuals with ocular anomalies, while heterozygous relatives remain asymptomatic (Plaisancié et al. 2018; Reis et al. 2021). This led us to investigate a potential maternally inherited non-coding variant affecting *FOXE3*.

Array-CGH (44k) analysis was performed to detect potential rearrangements near *FOXE3* that could impact its expression, but no such rearrangements were found. Sequencing of an approximately 600 bp evolutionarily conserved region, ∼3 kb upstream of *FOXE3* (GRCh37/hg19 chr1:47,878,072-47,878,696), revealed a rare heterozygous G>A variant at chr1:47,878,428 (rs745674596; with C as a possible alternative nucleotide) shared by the proband, her mother, and her sister (**Figures 1A and B**). The ultra-rare G>A substitution (with a minor allele frequency of 0.0001 in GnomAD v4.1, only observed in the heterozygous state) occurs at a nucleotide with high pathogenicity scores, including GERP (3.5799) and CADD GRCh37-v1.7 (18.83), indicating a potential deleterious effect. Notably, this variant is positioned 16 bp downstream of a 22 bp sequence, the deletion of which has been associated with complex microphthalmia and cataracts in mice (Wada et al. 2011) (**Figure 1B**). In addition, analysis of the variant using FATHMM-XF (Rogers et al. 2018), a predictive tool for evaluating the functional consequences of non-coding SNVs, yielded a non-coding score of 0.519910, strongly suggesting the variant’s pathogenicity. These findings collectively support a potential role of this non-coding variant in the proband’s disease. A homozygous G>C change at chr1:47,878,626 (rs10399673), inherited from both heterozygous parents, was also identified. This variant, with a minor allele frequency (MAF) of 0.444 in GnomAD v4.1, is likely benign. Whole Genome Sequencing (WGS) was subsequently performed to rule out other potential disease causes. It did not identify any additional candidate variants or rearrangements (including copy number and structural variants) in non-coding regions within *FOXE3* and its surrounding areas (**Figure S1**), especially in the topologically associating domain (TAD) regulatory unit comprising the gene (**Figure S2**). Additionally, after analyzing the dataset for both recessive and dominant inheritance patterns (the latter including *de novo* mutations, parental mosaicism, and incomplete penetrance), no other pathogenic variations were found that could explain the patient’s ocular phenotype.

### *In Vitro* Luciferase Assay indicates that the *FOXE3* rs745674596 Non-Coding G>A change acts as a regulatory variant

To assess whether the G>A change at position chr1:47,878,428 (rs745674596) affects *FOXE3* expression, we cloned into a BglI-digested pGL4.24 expression plasmid (Promega, USA), two fragments of 81 bp encompassing the rs745674596 G and A alleles (**Table S1**) upstream of a luciferase reporter gene for functional analysis. Luciferase assays revealed a 45-fold (p<0.0001) and 10-fold (not statistically significative p=0.2338) increase in expression for the mutant and wild-type constructs, respectively, compared to the empty vector (**Figure S3**). The 4.5-fold higher expression with the rs745674596 non-coding G>A mutant compared to the wild-type sequence suggests that this change should be classified as a regulatory variant (rv).

### Mice with the regulatory variant, truncating mutation, or both show genotype-correlated increased eye anomalies

To assess the impact of the rs745674596 non-coding G>A variation (rv) *in vivo*, we used CRISPR/Cas9 to edit the C57BL/6J mouse genome, generating mice with single heterozygosity, homozygosity, and compound heterozygosity for this regulatory variant and a premature termination codon (p.Gly28Argfs*112) resulting in a null (-) allele (**Figures 3A and 3C)**.

The resulting genotypes (*Foxe3+/rv*, *Foxe3+/-*, *Foxe3rv/rv*, *Foxe3rv/-* and *Foxe3-/-*) and wild-type controls (*Foxe3+/+*) were evaluated and compared for ocular globe size and structure using caliper measurements, slit lamp (**Table S4**) and OCT examinations. The *Foxe3+/+* mice (n = 30) exhibited no ocular anomalies (**Figure 2A and Figure S4 and S5**), with anteroposterior ocular diameters (APOD) ranging from 3.5 to 4 mm at three months of age, thereby establishing the normal range. The APOD of heterozygous *Foxe3+/rv* (28 animals, 56 eyes) and *Foxe3+/-* (22 animals, 44 eyes) mice remained within the normal range. However, some of these mice exhibited cataracts and anterior segment anomalies (**Figure 2A; Figure S4, S5 and Table S4**). The prevalence of ocular anomalies increased more strikingly in animals with these variants in homozygosity or compound heterozygosity, though their APOD also remained within the normal range. Specifically, 36% of *Foxe3rv/rv* (28 animals, 56 eyes) and 53% of *Foxe3rv/-* (40 animals, 80 eyes) mice exhibited cataract and anterior segment anomalies (corneal opacities, lens and iridocorneal adhesions, iris coloboma) (**Figures 2A, S4 and S5 and Table S4**). *Foxe3-/-* null mice (12/24 animals/eyes) exhibited a more severe and consistent phenotype, characterized by microphthalmia (APOD ≤ 3 mm) with significant anterior segment anomalies, including corneal opacities, central pits, iridocorneal and lens adhesions, athalamia, cataracts, and microphakia. This distinctive phenotype, present in 100% of *Foxe3-/- null* mice, aligns with the diagnosis of complex microphthalmia (**Figures 2A, S4 and S5 and Table S4**). Many of these mice also showed a small pyramidal pigmentary and vascular formation, suggestive of persistent fetal vasculature (PFV). Of note, the retina and its layers remained unchanged in most of the cases.

**Figure 2.**
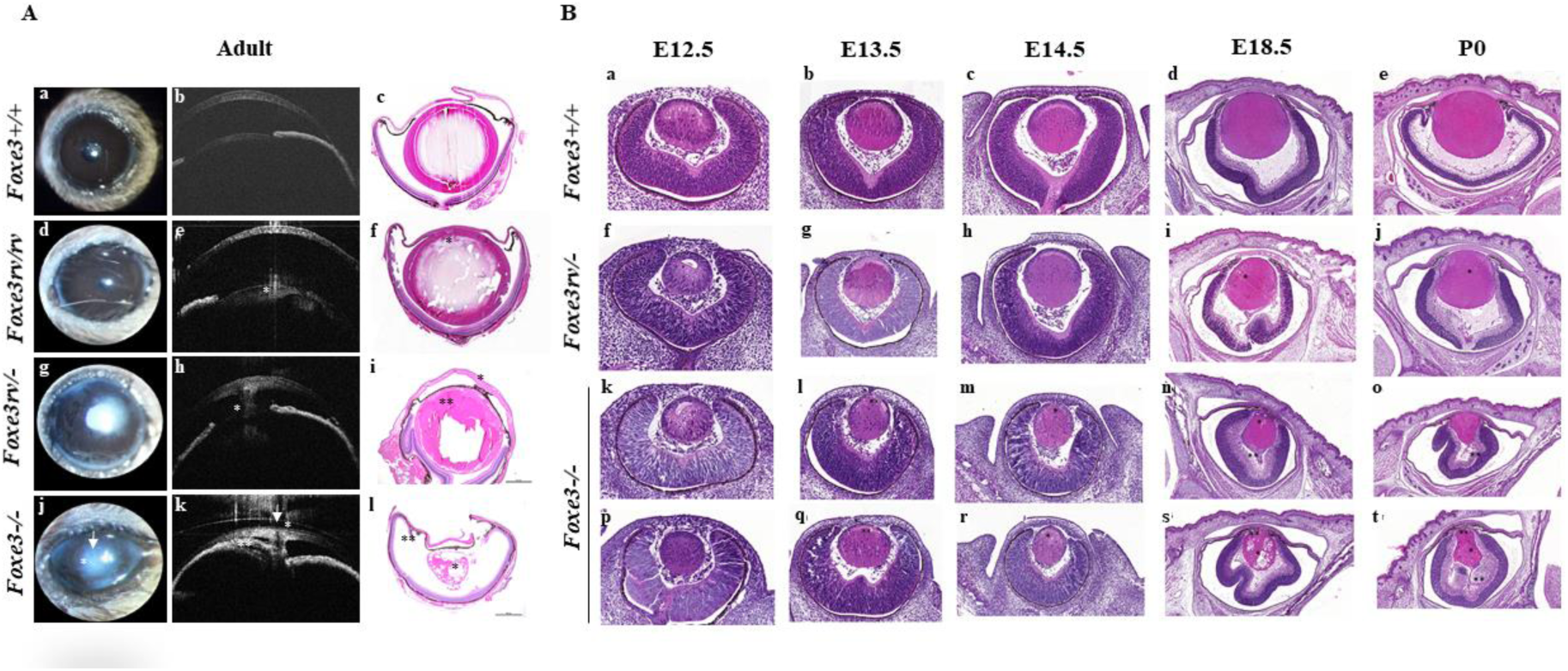
Comparative Histological Analysis of Ocular Development in *Foxe3* Mutants and Control Samples. (A) Representative live view, OCT images and histological sections and from adult wild-type mice (**a-c**) and mutant mice (n ≈ 25 per genotype) illustrate anomalies of increasing severity from focal anterior subcapsular cataract lesion in *Foxe3rv/rv* eyes (*)(**d-f**), to cataract (**)(**i**) with extensive irido-lenticular and irido-corneal adhesions and focal fibrotic thickening of the anterior lens capsule (*) (**h**,**i**) in *Foxe3*rv/- (**g-i**), to complex microphthalmia, with corneal clouding (*) (**j,k**), central pit (arrow) (**j,k**) and athalamia (**)(**j,k**), extensive uveo-corneal adhesions (**)(**l**), and a small, vacuolated, triangular-shaped cataractic lens (*) (**l**) in *Foxe3*–/–. (B) H&E-stained eye sections of *Foxe3*+/+, *Foxe3*rv/-, and *Foxe3-/-* mouse embryos from E12.5 to birth (P0) illustrate ocular development (**a-t**). **(a–e)** In *Foxe3*+/+ animals, early lens development is seen at E12.5, with the lens vesicle detaching from the surface ectoderm, a defined optic cup, and a neuroepithelial layer forming in the retina (**a**). By E13.5, the lens vesicle is rounder, with early fiber cell differentiation and thickening retina, marking early stratification (**b**). At E14.5, primary lens fibers elongate, the retinal ganglion cell layer becomes visible, and the optic nerve head connection develops (**c**). By E18.5, the lens and retina have mature features, including distinct retinal layers and defined anterior segments like the cornea and ciliary body (**d**). At P0, the lens is fully mature, with organized fiber cells and a defined capsule, while anterior structures like the cornea and iris continue developing (**e**). **(f–j)** In *Foxe3*rv/- animals, ocular development appears largely normal, with minor lens fiber vacuolization (*) (**i,j**), which may contribute to the adult cataract phenotype observed in non-null mice (**Table 2**). In some individuals, a small delay in lens detachment could be observed (*) (**g**). **(k–t)** In *Foxe3-/-* animals, initial development appears normal, but by E13.5, the anterior epithelial layer of the lens is disorganized (**) (**l,q**) and a delay in lens detachment is observed manifesting by a persistent lenticulo-corneal connection (*) (**l,q**). Mild vacuolization and swelling of lens fibers become apparent by E14.5, worsening over time (* in **m, r, s**, and **t**). The lens remains unusually close to the presumptive cornea, giving the appearance of an open lens at the anterior pole (** in **s** and **t**), with possible protein release into the corneal mesenchyme. The anterior epithelial layer gradually disappears, resulting in a microphakic lens (**n, o, s, t**). Additionally, inconsistent fibrosis is observed within the primary vitreous vascularization, extending from the posterior lens pole to the retina (**) (**n, o, t**).

We assessed the statistical significance of ocular defect penetrance (ranging from cataracts, anterior segment anomalies to complex microphthalmia), based on observed frequencies of 0% in *Foxe3+/+*, 18% in *Foxe3rv/+*, 23% in *Foxe3+/-*, 36% in *Foxe3rv/rv*, 53% in *Foxe3rv/-* and 100% in *Foxe3-/-* (**Figure S5**). Ocular penetrance in mice was significantly correlated with the number and the nature of the mutations **(Table 3)**. In the same manner, when classifying the phenotype in class of severity **(Table 1)**, the severity of the ocular phenotype was significantly associated with number and the nature of the mutations **(Table 3)**.

**Table 1.** Univariate analysis of potential association of the phenotype with age and/or gender. Age (in months) was significantly associated with the presence of a phenotype (p < 10⁻⁴), while gender showed no significant association; therefore, age but not gender was included in the logistic regression model with Firth’s correction, using *Foxe3+/+* as the reference genotype.

| Variable | OR | 95% CI | p-value |
| --- | --- | --- | --- |
| Intercept | 0.01 | 0.00-0.06 | 4.6<sup>e</sup>-07 |
| Age | 1;19 | 0.97-1.47 | 0.1 |
| Genotype |  |  |  |
| *Foxe3* +/+ | - | - | - |
| *Foxe3* rv/+ | 15.9 | 1.65-2131 | 0.013 |
| *Foxe3* +/- | 19.6 | 2.01-2629 | 0.007 |
| *Foxe3* rv/rv | 22.9 | 2.5-3053 | 0.002 |
| *Foxe3* rv/- | 63 | 7.81-8181 | 5.2<sup>e</sup>-07 |
| *Foxe3* -/- | 1305 | 60.4-413122 | 2.6<sup>e</sup>-10 |

**Table 2.**
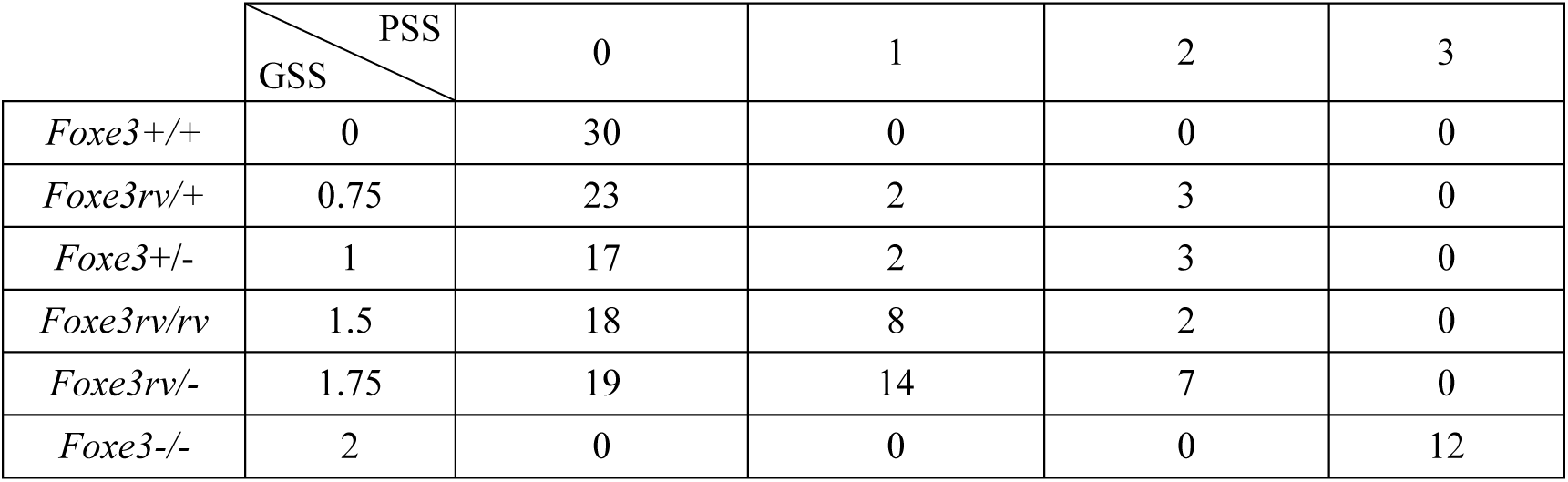
Distribution of analyzed mice by genotype across varying classes of genotypic and clinical severity. The scoring system quantifies genetic contributions to phenotypic outcomes. GSS (Genotype Severity Score): Sum of scores from both alleles, calculated from allelic scores of + = 0, rv = 0.75, and - = 1, representing overall genotype severity. PSS (Phenotype Severity Score): 0: No phenotype; 1: Mild; 2: Moderate; 3: Severe.

|  | PSS<br>GSS | 0 | 1 | 2 | 3 |
| --- | --- | --- | --- | --- | --- |
| *Foxe3*+/+ | 0 | 30 | 0 | 0 | 0 |
| *Foxe3*rv/+ | 0.75 | 23 | 2 | 3 | 0 |
| *Foxe3*+/- | 1 | 17 | 2 | 3 | 0 |
| *Foxe3*rv/rv | 1.5 | 18 | 8 | 2 | 0 |
| *Foxe3*rv/- | 1.75 | 19 | 14 | 7 | 0 |
| *Foxe3*-/- | 2 | 0 | 0 | 0 | 12 |

**Table 3.**
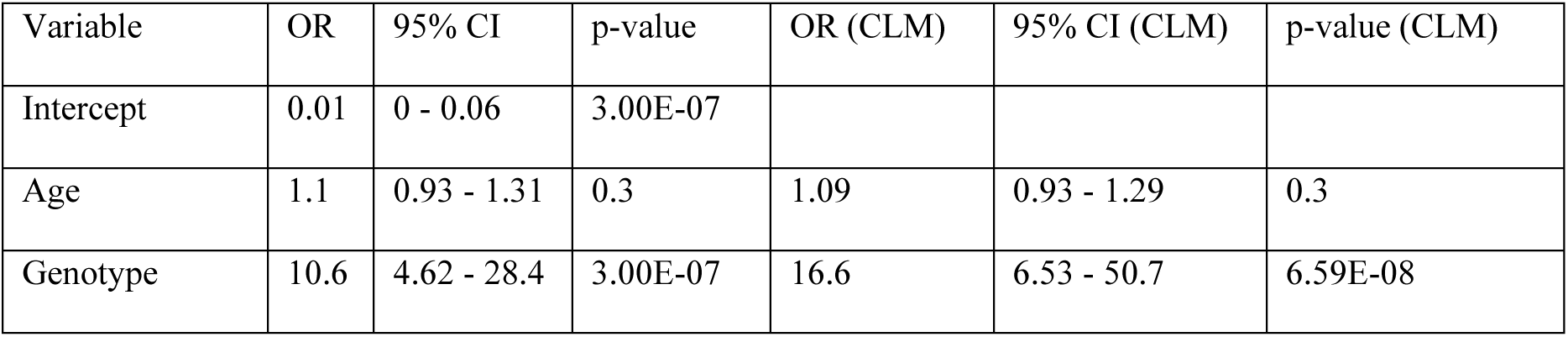
Correlation between Mutated Alleles and Phenotype Severity: Analysis of Odds Ratios (OR) from Statistical Regression Models. Analysis of the correlation between the number and severity of mutated alleles and the presence of a phenotype reveals statistically significant results, with ordinal regression analysis using cumulative link models (CLMs) demonstrating a significant association between the nature and severity of the mutated alleles and the severity of the phenotype (p < 0.01).

### Analysis of Ocular Development in *Foxe3* Mutant Mice Reveals Critical Role of FOXE3 in Lens Differentiation and Microphthalmia

To investigate the mechanisms behind *Foxe3*-related ocular anomalies, we compared ocular structures from compound heterozygous *Foxe3rv/-*, homozygous *Foxe3-/-*, and wild-type *Foxe3+/+* mice at key embryonic stages (E12.5, E13.5, E14.5, E18.5, P0). Histological analysis (3 animals, 6 eyes per genotype) showed nearly normal ocular development in *Foxe3 rv/-* animals, with slight lens fiber vacuolization potentially linked to cataracts (**Figure 2B**), as around 50% are expected to present this condition. In *Foxe3-/-* embryos, which all develop complex microphthalmia in adulthood, early lens structures were intact, but disorganization in the anterior lens epithelium and persistent lenticulo-corneal connections emerged by E13.5, leading to progressive lens abnormalities and degeneration (**Figure 2B**). These findings suggest that while *Foxe3* is expressed from E9.5 in the lens placode, it is not required for lens formation, as *Foxe3-/-* embryos had normal lens structure at E12.5 and *Foxe3* expression is confined to the anterior lens vesicle. However, *Foxe3* is critical for anterior epithelial cell elongation and differentiation into lens fibers starting at E12.5. Disruption of these processes leads to early and progressive lens abnormalities, contributing to the complex microphthalmia phenotype (**Figure 2B**).

### FOXE3 levels in the lens exhibit a genotype-dependent correlation in mice

To explore the correlation between *Foxe3* expression levels and genotypes, we measured *Foxe3* mRNA using real-time qPCR analysis from lens capsules of six adult animals per genotype, except for *Foxe3-/-* mice, whose microphakic eyes prevented proper sampling of the lens capsule. The capsules from both lenses of each animal were pooled and analyzed together. This study demonstrated a 50% reduction in *Foxe3* abundance (p<0.01) in the lens capsule and by inference in the anterior lens epithelium, the only tissue expressing *Foxe3* in the adult lens in *Foxe3rv/rv* mice compared to *Foxe3+/+* wild-type littermates (**Figure 3A**). Correspondingly, the *Foxe3* levels in heterozygous *Foxe3+/rv* mice were intermediate between those of *Foxe3rv/rv* and *Foxe3+/+* capsules (**Figure 3A**). Interestingly, contrary to the expected reduction in *Foxe3* abundance in *Foxe3+/-* and *Foxe3rv/-* mice, we observed a trend toward increased *Foxe3* levels in both genotypes compared to wild-type *Foxe3+/+* controls (**Figure 3A**). This was further supported by quantitative mRNA detection through RNAscope analysis of E12.5 embryos, which revealed significantly increased *Foxe3* staining in the anterior lens epithelium of *Foxe3-/-* null mice and decreased staining in *Foxe3rv/rv* mice compared to wild-type controls. In *Foxe3rv/-* mice, staining showed intermediate expression (**Figure 3B**), reflecting reduced expression from the allele carrying the regulatory variant and increased expression from the null allele, suggesting *Foxe3* upregulation associated with the p.Gly28Argfs*112 variation. FOXE3 protein levels were quantified through Western blot analysis, using total protein extracts from the lens capsules of each animal (three mice per genotype). The genotypes included mice with the regulatory variant in single heterozygosity, homozygosity, or compound heterozygosity with the null allele, as well as from the microphakic lenses of mice homozygous for the null allele. This analysis revealed a stepwise decrease in FOXE3 abundance across genotypes, from *Foxe3+/+* to *Foxe3+/-* to *Foxe3rv/rv* to *Foxe3rv/-* to *Foxe3-/-* (**Figure 3C**). Notably, despite increased mRNA levels, FOXE3 protein abundance was markedly reduced in lenses with the null allele, with a decrease of approximately 50% in *Foxe3+/-* mice and around 75% in *Foxe3rv/-* mice, and FOXE3 was virtually undetectable in *Foxe3-/-* lenses (**Figure 3C**).

**Figure 3.**
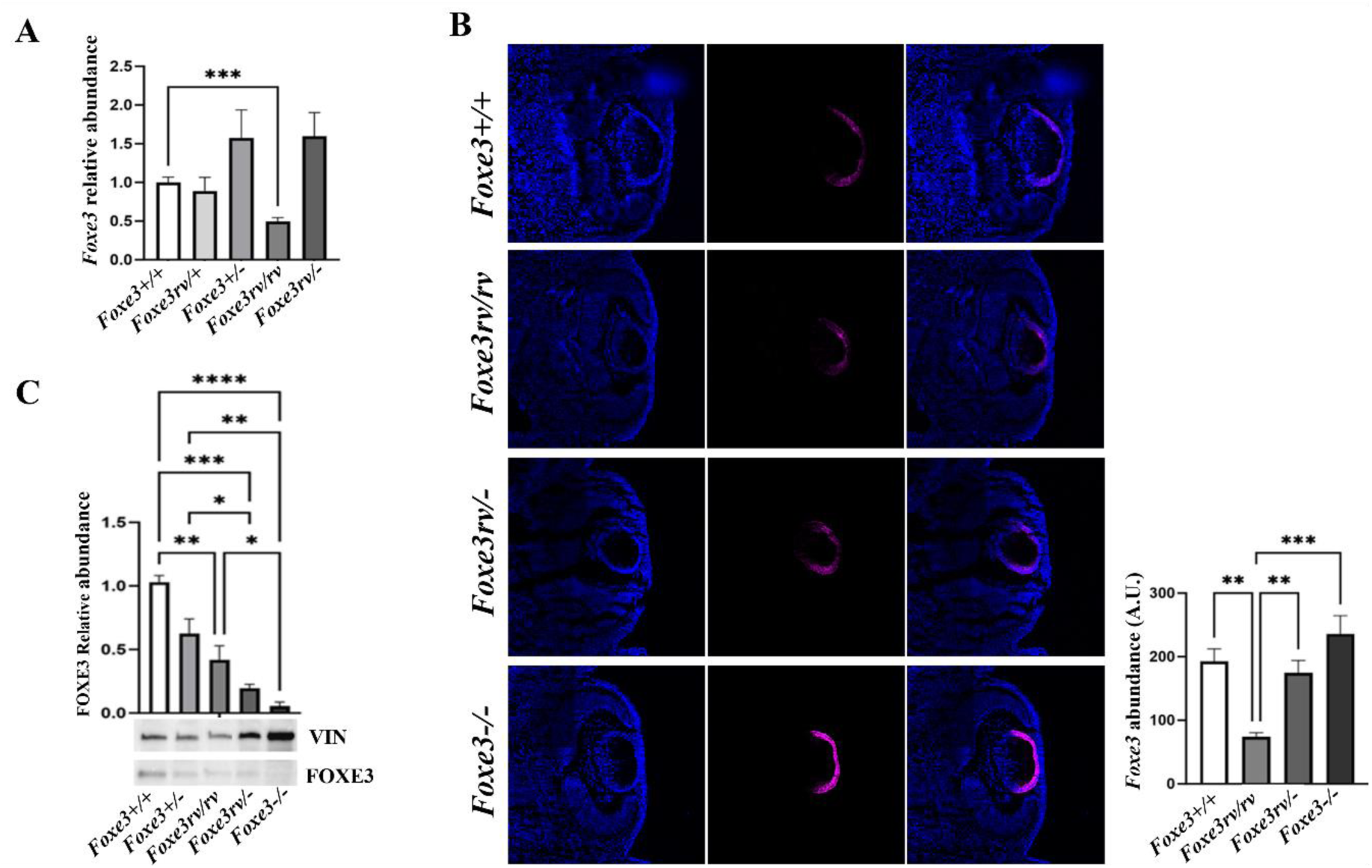
Comparative Analysis of *Foxe3* mRNA and Protein Expression in Lens Epithelium of *Foxe3* Mutant and Control Mice. (A) Relative *Foxe3* abundance in the anterior lens epithelium was measured by RT-qPCR in *Foxe3*+/+, *Foxe3*rv/+, *Foxe3*+/-, *Foxe3*rv/rv, and *Foxe3*rv/- mice. (B) RNAscope *in situ* hybridization provides quantitative mRNA detection in tissue sections; however, the probes cannot differentiate between wild-type *Foxe3* mRNA and the frameshift variant produced in -/- mice. *In situ* hybridization was performed on lens sections from *Foxe3*+/+, *Foxe3*rv/rv, *Foxe3*rv/-, and *Foxe3*–/– mice at E12.5. (C) FOXE3 protein levels in the lens epithelium, as determined by Western Blot (WB). Quantitative analysis of WB signals shows a significant reduction in FOXE3 protein abundance with the rv variant. Only statistically significant differences are indicated.

Together, these results provide strong support to the pathogenicity of the rs745674596 non-coding G>A variant through decreasing expression of the gene and underscore a correlation between reduced FOXE3 protein levels and an increase in the penetrance and severity of ocular anomalies.

### DNA Pull-Down and Mass Spectrometry Reveal Interactions Influencing *FOXE3* Expression by the rs745674596 Non-Coding G>A Variant

DNA pull-down and mass spectrometry analyses were performed to determine whether the rs745674596 non-coding G>A variant influences *FOXE3* expression by affecting the binding of specific transcription factors. We utilized whole lysates from immortalized HLEpiC and nuclear extracts from mouse CCE-RAX and CCE cell lines, using oligonucleotides specific to either wild-type or mutant human non-coding sequences (**Table S1**).

Our experiments identified 16, 16, and 17 transcription factors differentially bound to wild-type and mutant oligonucleotides in HLEpiC, CCE-RAX, and CCE cell lines, respectively (**Figure 4A**). Notably, USF2, GABPA and GABPB1 were common across all three cell lines, though GABPB1 was excluded from further analysis due to its role as a GABPA cofactor. CNBP, linked to microphthalmia in mice,^30^ was also considered despite being detected only in HLEpiC and CCE cells. GABPA demonstrated a preference for binding to the mutant variant over its wild-type counterpart. In contrast, USF2 and CNBP displayed stronger binding to the wild-type version, suggesting that their interactions were either partially or completely diminished in the mutant form.

**Figure 4.**
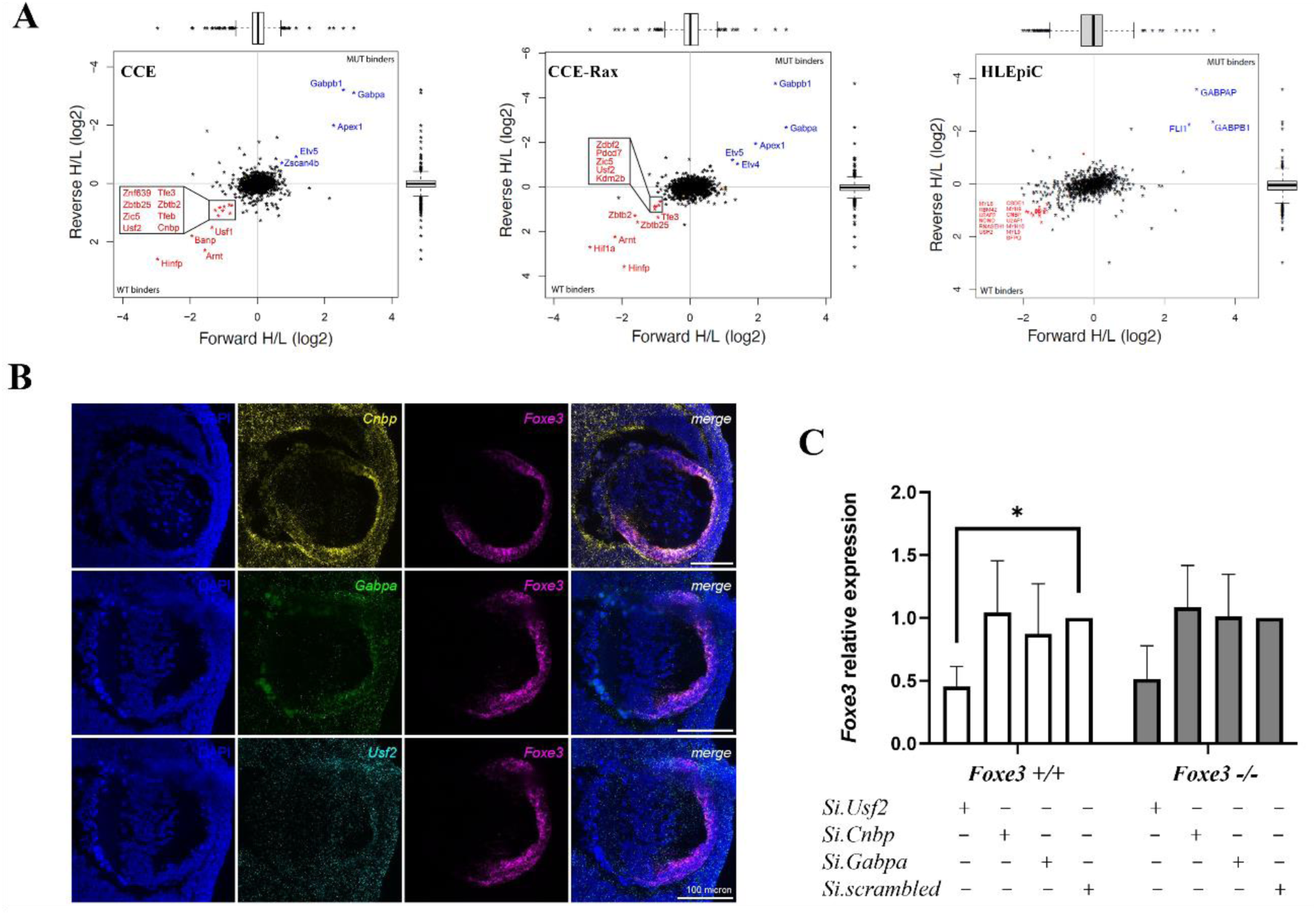
Identification and Functional Analysis of Transcription Factors Binding to *Foxe3* Regulatory Element in Lens Cells. (A) Combined DNA-PD and mass spectrometry analysis in murine CCE cells, CCE-Rx cells, and immortalized HLEpiC. Factors with a significant preference for binding the wild-type oligonucleotide are marked in red (p<0.05), while those preferring the mutant oligonucleotide are marked in blue (p<0.05). (B) RNAscope in situ hybridization shows spatiotemporal expression patterns of candidate genes. At E12.5, *Foxe3* expression localizes within the anterior layer of the presumptive lens, while *Cnbp*, *Gabpa*, and *Usf2* display ubiquitous expression. (C) Relative *Foxe3* expression in lens epithelial cells from *Foxe3+/+* and *rv/rv* genotypes after inhibition of *Cnbp*, *Usf2*, and *Gabpa*, as measured by RT-qPCR. (* p<0.05).

RNAscope *in situ* hybridization of developing mouse eyes demonstrated colocalization of *Usf2*, *Cnbp*, and *Gabpa* with *Foxe3* mRNA in the anterior lens epithelium, though their expression was broader compared to the lens-specific expression of *Foxe3* (**Figure 4B**).

Knockdown experiments using siRNAs revealed a statistically significant reduction in *Foxe3* mRNA in *Foxe3+/+* cells (p = 0.0322) and a marked trend towards reduction in *Foxe3rv/rv* cells upon *Usf2* inhibition. In contrast, *Cnbp* and *Gabpa* depletion did not affect *Foxe3* expression (**Figure 4C**).

These results suggest that the rs745674596 non-coding G>A variant impacts *FOXE3* expression by reducing *Foxe3* levels through its effect on USF2 binding, thereby further supporting the pathogenicity of the variant and highlighting the critical role of USF2 in lens and eye development.

## DISCUSSION

The *FOXE3* region, located within a 50 kb TAD on chromosome 1p33, contains several evolutionarily conserved sequences upstream of the gene. Among these, the chr1:47877964-47878774 region (GRCh37/hg19), located 3 kb upstream of *FOXE3*, has been linked to reduced *Foxe3* expression and mild microphthalmia with cataracts in the rct mutant SJL/J mouse carrying a 22-bp homozygous deletion (Wada et al. 2011). Consistent with a regulatory role, this region displays open chromatin and DNase hypersensitive areas, despite lacking CpG islands. Studies have also shown binding of transcription factors such as SOX2, PAX6, PITX3, and TFAP2A, which are crucial for ocular development (Zhao et al. 2019). Therefore, genetic alterations in this conserved region have the potential to impact *FOXE3* expression, leading to either downregulation or upregulation of *FOXE3* and contributing to both recessive and dominant *FOXE3*-associated diseases reported in the literature.

Conforming with established genotype-phenotype correlations linking recessive *FOXE3* mutations to the most severe *FOXE3*-related diseases (Plaisancié et al. 2018), the ultra-rare non-coding rs745674596 G>A change at chr1:47878428 identified in *trans* of the *FOXE3* c.720C>A (p.Cys240*) truncating mutation in a patient with complex microphthalmia, was expected to result in a loss-of-function through the dampening of *FOXE3* expression. Intriguingly, luciferase reporter assay analysis of the variant revealed increased transactivation activity compared to the wild-type. While supporting an impact of the change on *FOXE3* expression, this observation challenged the recessive inheritance model and suggested the possibility of dominant inheritance with incomplete penetrance. However, in contrast to the *in vitro* results, *in-vivo* studies revealed markedly reduced *Foxe3* expression in the lens of mice homozygous for non-coding rs745674596 G>A variant, confirming the expected loss-of-function mechanism. This discrepancy highlights the limitations of basic *in-vitro* analyses, such as luciferase reporter assays, in assessing pathogenic mechanisms of non-coding variants. Regulatory regions can contain both activating and repressing motifs, and gene expression regulation can vary over time and across different tissues. Another concern regarding mRNA level analysis arises from the unexpectedly elevated levels of *Foxe3* mRNA with the p.Gly28Argfs*112 frameshift mutation observed in the developing lenses of *Foxe3-/-* mice compared to wild-type littermates. This issue is further highlighted by comparable *Foxe3* mRNA levels in wild-type and *Foxe3rv/-* mice, where reduced expression from the non-coding rs745674596 G>A allele is masked by increased expression of the frameshift mutation. The mechanisms behind elevated levels of mRNA encoding premature termination codons remain unclear. Given that *Foxe3* is a single-exon gene and both null and wild-type mice share the same genetic background, splicing and increased transcription are unlikely to explain the higher levels of mRNA encoding the p.Gly28Argfs*112 frameshift variation. This accumulation is more likely driven by the absence of Nonsense-Mediated mRNA Decay (NMD) or the stabilization conferred by RNA-binding proteins or other factors that prevent degradation (Millevoi and Vagner 2010). Whether the c.720C>A nonsense mutation (p.Cys240*) identified in the individual has a similar impact on *FOXE3* content in the lens remains an open question. Yet, the evidence that both the non-coding variant and the frameshift mutation had unexpected effects on expression levels underscores the pitfalls of relying solely on RNA levels as biomarkers for pathogenicity.

In contrast to mRNA studies, which failed to link *Foxe3* expression with the penetrance of ocular defects, analysis of FOXE3 protein content in the lens of mice revealed a correlation between FOXE3 levels and both the frequency and severity of ocular anomalies. The spectrum of ocular anomalies affecting eye growth, cornea, anterior segment and lens, along with their graded frequency and severity in relation to *Foxe3* expression in mice, aligns with human findings, where genotype-phenotype studies have linked the penetrance of these anomalies to the number and type of *FOXE3* mutations (Plaisancié et al. 2018; Reis et al. 2021).

More specifically, the phenotype in our *Foxe3-/-* null mice exhibited normal eye structure by E12.5. However, from E13.5 to E14.5, mild and discrete abnormalities emerged, including delayed lens vesicle detachment, disorganized anterior lens epithelium, mild vacuolization, and abnormal lens positioning, resulting in microphakia by E18.5. In adults, we observed a consistent phenotype of complex microphthalmia in all mice, accompanied by severe anterior chamber lesions and lens anomalies, but no "true" anophthalmia. These findings align with the established role of the FOXE3 transcription factor in lens epithelial cell growth, where it regulates proliferation, apoptosis, and cell cycle (Wang et al. 2012; Islam et al. 2015). *Foxe3* expression begins in the lens placode at E9.5, facilitating lens vesicle closure and separation from the ectoderm. This expression continues through the lens vesicle stage at E12.5, playing a crucial role in balancing the proliferation and differentiation of lens fiber cells and the anterior lens epithelium into adulthood (Blixt et al. 2007).

The observed findings are consistent with other *Foxe3* mutant lines, including the homozygous dyl/dyl mutant (Blixt et al. 2007; Sanyal and Hawkins 1979) and engineered null mutants from 129×C57Bl/6 (Medina-Martinez et al. 2005) and Balb/c mice (Blixt et al. 2007), which also display severe lens abnormalities and eye growth defects such as microphthalmia. A common phenotype among these mutants is a persistent connection between the lens and cornea due to delayed lens vesicle closure, resulting in lens with reduced size, irregular shape and structural disorganization with vacuoles. Although fiber elongation and crystallin expression initiate normally, the number of fibers drastically declines, leading to cataracts and, in some cases, fiber expulsion through the ectodermal connection. Both dyl/dyl and null mutants show impaired lens epithelium proliferation and failed lens fiber differentiation (Blixt et al. 2007; Medina-Martinez et al. 2005). Notably, the E14.5 anterior epithelium in mutants displays many apoptotic cells, unlike wild-type lenses. Similar findings in rct/rct mice (Wada et al. 2011; Maeda et al. 2001), which carry a 22 bp deletion in the *Foxe3* regulatory region, demonstrate mild microphthalmia and significant lens degeneration.

The first phenotypic deviation in *Foxe3* mutants is the delayed separation of the lens vesicle from the ectoderm, and secondarily, anterior lens epithelium disorganization, leading to lens degeneration. However, the mechanism linking this degeneration to microphthalmia remains unclear. Possible explanations include the larger size of the lens in mice compared to humans, suggesting that its absence could collapse the eye and cause microphthalmia; however, cases of aphakia without microphthalmia have been observed in humans (Reis et al. 2021). The persistent ectodermal connection may reduce intraocular pressure, limiting eye growth, yet this connection is also found in some compound heterozygous *Foxe3rv/-* individuals without microphthalmia (**Figure 2B, g**). The *Foxe3* mutant model resembles cavefish (Astyanax mexicanus), where early lens apoptosis leads to complete eye degeneration, leaving a non-functional cyst covered by skin in adults (Rétaux 2022; Alunni et al. 2007; Yamamoto and Jeffery 2000). Transplant experiments reveal the lens’s role in degeneration: a cavefish lens induces degeneration in surface embryos, while a surface lens can rescue cavefish eyes (Yamamoto and Jeffery 2000). Although alpha-A crystallin deregulation may play a role (Hinaux et al. 2015), the genetic basis for lens apoptosis in that species remains unknown.

In all vertebrates, optic vesicle folding coincides with lens vesicle formation, and their interaction is essential for proper eye morphogenesis (Casey et al. 2023). Lens-retina crosstalk, mediated by factors and pathways (e.g., BMP, FGF, Wnt), is critical for eye development (Casey et al. 2023). Studies have explored interactions among the lens placode and optic vesicle, the lens vesicle and retina, as well as periocular mesenchyme and neural crest cells. PAX6 is essential for lens placode development but not for optic vesicle induction (Collinson et al. 2000; Davis-Silberman et al. 2005; Dimanlig et al. 2001). The "induction" hypothesis by Spemann H (SPEMANN 1901) is being reevaluated, with evidence showing optic cup formation can occur without lens induction (Cardozo et al. 2023). The presence of optic cups in aphakic patients further supports this (Valleix et al. 2006). The lens, evolving secondarily, may primarily enhance optic cup invagination and formation (Cardozo et al. 2023). Our study proposes a model of eye growth defects wherein initial developmental cues successfully establish ocular structure formation. However, as development advances, "inductive" processes are supplanted by "involutive" ones, culminating in lens degeneration and disrupted ocular growth. The Aey69 mutant line (Vetrivel et al. 2019) also displays microphthalmia, aphakia, and retinal hyperproliferation due to a *Hist2h3c1* mutation, disrupting lens development and affecting gene expression in retinal progenitor cells. This raises the question of whether complete eye involution (as in cavefish) and incomplete involution with retinal hyperproliferation (other mouse models) involve the same or distinct mechanisms leading to eyeball degeneration.

Due to challenges in accessing and culturing embryonic lens epithelial cells, we selected the most relevant publicly available cell types to investigate transcription factor binding to the non-coding region surrounding the non-coding rs745674596 G>A allele variant. We selected CCE mouse embryonic stem cells for their broad gene expression profile and CCE-RAX for their relevance to eye development, given that RAX is essential for the formation of the retina and other ocular structures during embryogenesis. Additionally, we immortalized human epithelial lens cell line to ensure our study encompassed relevant human lens biology. DNA-PD and mass spectrometry using these cell lines revealed a limited set of 34 candidate transcription factors, which displayed differential binding in the presence and absence of the variant. To enhance selectivity, we focused on transcription factors shared among the lines or known to play a role in eye development, resulting in a shortlist of three candidates. Notably, none of them were identified through *in-silico* analysis using CHIP-seq data from the ENCODE project or JASPAR predicted transcription factor binding sites. SiRNA knockdown analysis indicated that only USF2 influenced *Foxe3* expression. The other two candidates, particularly CNBP - associated with microphthalmia in mice (Chen et al. 2003) - may still play a role in lens development *in vivo*, especially given their expression patterns during ocular development. Both colocalize with *Foxe3* in the lens vesicle during early stages and later in the lens epithelium, suggesting a potential overlap in function. However, further investigations are needed.

*Foxe3* expression decreased upon *Usf2* inhibition in both wild-type and rv/rv cells, with statistical significance achieved in wild-type but not in rv/rv. This finding aligns with DNA pull-down assay results, which showed preferential binding of USF2 to the wild-type sequence compared to the rv variant. The *USF2* gene encodes a transcription factor from the basic helix-loop-helix leucine zipper (bHLH-LZ) family (Huang et al. 2023). This protein plays a role in facilitating transcription by interacting with pyrimidine-rich initiator (Inr) elements (Breen and Jordan 1999) and E-box motifs (CACGTC) (Xie and Herschman 1996; Vivas-Mejía et al. 2009). Interestingly, the rs745674596 G>A variant specifically alters the E-box motif within the *FOXE3* regulatory region, potentially reducing the binding affinity of USF2 to the DNA.

USF2 plays a critical role in regulating cell proliferation, differentiation, and apoptosis, processes essential for tissue development and maintenance (Chi et al. 2020). *Usf2* knockout mouse model exhibits growth delay, apathy and increased postnatal lethality (Sirito et al. 1998; Vallet et al. 1997), though ocular malformations have not been specifically investigated. Interestingly, Fujimi and Aruga demonstrated *USF2* expression in neural tissues, including the eye, during key stages of embryonic development in Xenopus laevis (Fujimi and Aruga 2008). Our findings similarly show that *Usf2* is expressed in ocular structures during mouse development, indicating a possible link between USF2 and ocular anomalies. Given its role in the development of neural tissues, including the eye (Fujimi and Aruga 2008), disruptions in *USF2* expression or function may contribute to ocular developmental defects. Although no direct link between *USF2* mutations and human disease has been confirmed, exploring *USF2* variants in individuals with ocular anomalies could yield important insights. Further investigations may uncover critical associations between USF2 and eye development or related disorders. Several truncating mutations in *USF2* have been reported in large-scale genome databases (GnomAD), but only once at the homozygous state, providing support for its possible implication in human autosomal recessive diseases.

In conclusion, this study highlights the significance of highly conserved non-coding regions in understanding genetic diseases while emphasizing the challenges in interpreting variants within these regions. By focusing on genotype-phenotype correlations, we demonstrate their critical role in elucidating the impact of non-coding variants, particularly in single heterozygous cases of potential recessive diseases where biallelic mutations are absent, thus hampering optimal genetic counselling. Fully characterizing such variants requires a comprehensive approach that integrates in-depth knowledge of disease expression and genetics, along with multiple lines of experimental evidence of pathogenicity, rather than relying solely on standard molecular diagnostics. Furthermore, not only this study underscores the medical relevance of searching for pathogenic regulatory variations, it also highlights the impact of these analyses on advancing our understanding of gene regulation and identifying novel candidate genes for genetic diseases. Finally, we provide the first description of a novel mechanism causing microphthalmia via lens degeneration, which carries significant therapeutic implications. Previously, treatment options were limited due to early ocular developmental defects; however, the discovery of a degenerative mechanism opens new possibilities for intervention, especially if it occurs later in pregnancy.

## FAMILY, MATERIALS AND METHODS

### Clinical and genetic analysis

Toulouse University Hospital (TUH) Data Protection Officer has ensured this study complies with France CNIL MR-004 standards for medical research and adheres to EU General Data Protection Regulation requirements. The study was included in both the TUH retrospective study registry (RNIPH #2021-44) and the CNIL MR-004 registry (#2206723 v 0).

Full medical and familial history was collected. The patient underwent detailed general and ophthalmological examination with slit lamp examination, gonioscopy, ocular ultrasound, fundus examination and optical coherence tomography (OCT). Brain magnetic resonance imaging (MRI) was performed to investigate visual tract and intracerebral structures. Ultrasound was used to investigate cardiac and kidney structures.

The patient underwent analysis of 119 genes involved in ocular development using a customized NGS panel, as previously described (Chesneau et al. 2022). This method enables the detection of both single nucleotide variants (SNVs) and copy number variations (CNVs). Variants were classified according to American College of Medical Genetics and Genomics (ACMG) guidelines (Richards et al. 2015). Candidate variants and their co-segregation with the disease within the family **(Figure 1A)** were confirmed by Sanger sequencing with specific primers, available upon request.

CGH-array (BlueGnome, UK, 44k) was performed on the proband’s DNA to identify genomic rearrangements, following previously described protocols (Evangelidou et al. 2013).

Whole-genome sequencing (WGS) of the proband and her parents was performed by the Centre National de Recherche en Génomique Humaine (CNRGH) in Evry, France. Genomic DNA (1 µg) from each sample was used to prepare whole-genome sequencing libraries with the Illumina TruSeq DNA PCR-Free Library Preparation Kit (Illumina Inc.), following the manufacturer’s instructions. Libraries were sequenced on an Illumina NovaSeq 6000 platform as paired-end 150 bp reads, with samples pooled on a NovaSeq 6000 S4 flowcell to achieve a minimum average depth of 30x. After demultiplexing, sequences were aligned to the hg19 reference genome using the Burrows-Wheeler Aligner. Subsequent processing was done with GATK, SAMtools, and Picard following established best practices (http://www.broadinstitute.org/gatk/guide/topic?name=best-practices). Variants were called using GATK Haplotypecaller v4, and large variants (CNVs and SVs) were identified using WiseCondor (Raman et al. 2019), Canvas (Roller et al. 2016) and Manta(Chen et al. 2016).

All variants were filtered using an in-house developed annotation software system (Polyweb), based on two inheritance modes: recessive and dominant (including *de novo* mutations, parental mosaicism, and incomplete penetrance) with minor allele frequencies (MAF) of <1% and <0.1%, respectively. The pathogenicity of the selected variants was assessed using the Alamut Mutation Interpretation Software (http://www.interactive-biosoftware.com), a decision support system for mutation interpretation based on Align DGVD, MutationTaster, PolyPhen-2, SIFT, SpliceSiteFinder-like, MaxEntScan, NNSPLICE, GeneSplicer, Human Splicing Finder, ESEfinder, and RESCUE-ESE. The pathogenicity of the identified non-coding variant was assessed using scores such as GERP, CADD GRCh37-v1.7 and FATHMM-XF (Rogers et al. 2018).

### Hi-C data visualization

To characterize the chromosome conformation structure of the *FOXE3* locus in humans, we analyzed publicly available Hi-C data from the 4D Nucleome Data Coordination and Integration Center. This data, generated from H1-hESC cell lines, was processed using Micro-C and its updated version, Micro-C XL (Hsieh et al. 2016). These methods provide nucleosome-level resolution, with Micro-C XL offering improved recovery of higher-order interactions through enhanced cross-linking. Data from both methods were combined into contact matrices for each cell line, which were further processed into binary heatmap files, similar to the *.hic* files used by the UCSC Genome Browser.

For chromatin conformation analysis of the *Foxe3* locus in mice, we used mESC Hi-C data from Bonev et al. (Bonev et al. 2017), generated from intact cell nuclei using restriction enzyme-mediated DNA cleavage prior to proximity-based ligation (Rao et al. 2014). Hi-C reads were trimmed and quality-checked using Trim Galore (v0.6.5, Cutadapt v2.6, FastQC v0.11.9) (Krueger K. 2015). They were mapped to mm10 and filtered for Hi-C artifacts using HiCUP (v0.7.2, Bowtie2 v2.3.5, R v3.6.0_3.9) (Wingett et al. 2015). Contact maps were generated and analyzed with Juicer (v1.22.01) (Durand et al. 2016) using the parameter -s DpnII and resolution determined via the “calculate_map_resolution.sh” script. HOMER was used to compute observed matrices, adjusting for linear distance and sequencing depth. The resulting Hi-C contact maps were visualized as UCSC custom tracks on the UCSC Genome Browser.

### Luciferase Reporter Assay Analysis of rs745674596 G and A Alleles

Two complementary single-stranded 81-base oligonucleotide pairs spanning GRCh37/hg19 chr1:47,878,388-47,878,468 and representing the rs745674596 G (WT) and A (Mut) alleles, with three additional nucleotides at their 3′ ends to introduce BglI cohesive ends, were designed and synthesized by Sigma-Aldrich Technologies (**Table S1**). The WT and Mut oligonucleotide pairs were hybridized separately and then cloned into the BglI-digested pGL4.24 expression plasmid (Promega, USA), positioning them upstream of the PGK1 promoter and the firefly luciferase reporter gene. T4 DNA ligase in 1X reaction buffer (NEB, UK) was used according to the manufacturer’s protocol. The resulting WT and Mut pGL4.24 constructs were transformed into competent E. coli (OneShot TOP10, Thermo Fisher Scientific, Asnières-sur-Seine, France) and propagated to amplify the constructs. The plasmids were extracted and purified using a Miniprep kit (Qiagen, Hilden, Germany), according to the manufacturer protocol. The insertion and orientation of the mutant and wild-type inserts were confirmed by Sanger sequencing of the plasmids.

Undifferentiated CCE-Rx mouse embryonic stem cells (mESCs) expressing the Retina and Anterior neural fold homeoboX (*Rax*), a key gene in retina and lens formation (Tabata et al. 2004), were cultured in gelatin-coated (Biocoat, USA) 10 cm dishes with DMEM supplemented with pyruvate, glucose and Glutamax (Thermo Fisher Scientific), β-mercaptoethanol (1%), MEM non-essential amino acids (Thermo Fisher Scientific) (1%), Penicilin-Streptomycine, embryonic stem (ES) cell culture medium supplemented with fetal calf serum (FCS) (Dutscher) (15%), and leukemia Inhibitory Factor (LIF, Sigma-Aldrich). Cells were kept at 37°C and 5% CO₂, with daily medium changes and subculturing every 48 hours. For transfection, the cells were trypsinized and seeded in non-adherent dishes to promote embryoid body formation (1,2.10^6^ of cells in a 6 cm plate containing LIF-free medium) and incubated at 37°C / 5% de CO_2_. Trypsinized cells were co-transfected using a recombinant vector (1 µg of either unmodified pGL4.24 vector or the mutant or wild-type constructs), Renilla reporter vector (100 µg; 100:1 ratio), and enhancer DNA in Effectene reagent, following the manufacturer’s protocol (Qiagen, Hilden, Germany). Non-transfected CCE-Rx cells served as a negative control for background noise. Forty-eight hours post-transfection, embryoid bodies were harvested, washed with 1X PBS, and lysed with Passive Lysis Buffer (PLB) (Promega, USA). Lysates were stored at -20°C. Luciferase activity measurements were conducted in 96-well plates, with 20 µl per well, based on three independent transfections for each condition, with each transfection performed in triplicate. A luminometer (NOVOstar, BMG labtech) recorded absolute luminescence emitted by both Firefly and Renilla luciferases activities, and the Firefly/Renilla luminescence ratio was calculated for each sample. The normalized ratios were analyzed using a one-way ANOVA, followed by Tukey’s multiple comparisons test to identify significant differences between means (*** = p < 0.0001). Data are presented as mean ± standard deviation (SD) on the graph.

### Immortalization of Human lens epithelial cells

Human lens epithelial cells (HLEpiC; Innoprot, Derio, Bizkaia, Spain) were cultured in 6-well plates using Epithelial Cell Medium (EpiCM; Innoprot) supplemented with 10% fetal bovine serum (FBS) at 37°C in 5% CO₂. The cells were immortalized by transfection with 1 µg of the pLAS plasmid encoding the SV40 large T antigen (Daya-Grosjean et al. 1987), using Lipofectamine 2000 (Thermo Fisher Scientific) according to the manufacturer’s protocol. The cells were transferred to a T25 flask, propagated to confluence, and the SV40 large T antigen-expressing immortalized HLEpiC were isolated based on morphology through cell sorting into 96-well plates for subsequent expansion and experimentation.

### DNA pull-down and mass spectrometry

DNA pull-down experiments were conducted using nuclear extracts from undifferentiated CCE-Rx and CCE cells, along with whole cell lysates from immortalized HLEpiC. CCE-Rx and CCE cells were cultured as previously described, harvested by trypsinization, and washed with PBS. Nuclear extracts were then prepared from both cell types following established protocols (Karemaker and Vermeulen 2018). For whole cell lysate preparation, immortalized HLEpiC were cultured in EpiCM supplemented with 10% FBS, harvested by trypsinization, washed with PBS and then resuspended in a buffer containing 150 mM NaCl, 50 mM Tris (pH 8.0), EDTA-free complete protease inhibitors (CPI) (Roche), and 1% NP40.

The biotinylated 81 bp oligonucleotides encompassing the rs745674596 G and A alleles (FOXE3-WT_F and FOXE3-Mut_F sequences; see **Table S1**) were sourced from Integrated DNA Technologies (IDT, Leuven Belgium). DNA pull-down experiments were conducted using 150 µg of nuclear extract or 1.5 mg of whole cell lysate per reaction, following a previously described protocol (Karemaker and Vermeulen 2018). Proteins were digested on-bead into tryptic peptides, which were subsequently cleaned and eluted using StageTips. The light- and medium-labeled samples were then combined in both forward and reverse reactions. Peptides were then separated on an Easy-nLC 1000 (Thermo Fisher Scientific, Waltham, Massachusetts, US) connected online to an LTQ-Orbitrap Fusion Tribrid mass spectrometer (Thermo Fisher Scientific) or an Exploris 480 mass spectrometer (Thermo Fisher Scientific), using a 2h or 1h linear gradient of acetonitrile, respectively. For samples measured on the Fusion, scans were collected in data-dependent top-speed mode of a 3 s cycle with dynamic exclusion set at 60 s. For samples measured on the Exploris, scans were collected in data-dependent top-20 mode with a 45 s dynamic exclusion. Peptides were searched against the UniProt mouse or human proteome with MaxQuant v1.5.1.0 (Cox and Mann 2008), using default settings, the appropriate dimethyl labels, and re-quantify enabled. Data were analyzed with Perseus version 1.4.0.0 and R.

### Generation of Foxe3 mouse models

Mouse lines harboring the regulatory rs745674596G>A variant (*Foxe3^em1Goph^*allele, hereafter referred to as the *rv* allele) upstream of *Foxe3*, or a premature termination codon in the gene (c.82_92del; p.Gly28Argfs*112; *Foxe3^em2Goph^*, referred to as the *-* allele) were created using the CRISPR/Cas9 system by the Transgenesis platform at the LEAT Facility (Imagine Institute). For the *rv* allele, the guide RNA (sgRNA: 5’-ATGCTCAGCCGCATCACGTC-3’; designed with CRISPOR (http://crispor.tefor.net/)) and a phosphorothioate-modified ssODN (5’A*T*CCAGGCCCATGAGAAAGGGGCCACCTTCACTGGCCGTCTTATGCCCGGAT GCTCAGCCGCATCAC<u>A</u>TCCGGCCCAGGGCCTGTGAAAAGAGGGCCCAGCCACGCT GAAAACGCGGA*T*T-3’) centered on the rs745674596 variant (underlined) were used. For the - allele, the CRISPOR-designed sgRNA CTCCGGTTCGCGCCCCGGTT was employed. The CRISPR/Cas9 ribonucleoprotein (RNP) complex was microinjected into C57BL/6J mouse zygotes’ pronuclei, following the method of Ucuncu et al. (Ucuncu et al. 2020). Offspring were genotyped using PCR amplification and Sanger sequencing with appropriate primers **(Table S2)**. To minimize potential off-target mutations, mutant mice were backcrossed to C57BL/6J mice for several generations before intercrossing to obtain *Foxe3+/+, Foxe3rv/+*, *Foxe3rv/rv*, *Foxe3+/-, Foxe3-/- and Foxe3rv/-* lines. Animal procedures were approved by the French Ministry of Research and complied with the French Animal Care and Use Committee from Paris Descartes University (APAFIS# 31491).

### Ocular Phenotype Assessment in Adult Mice

Adult mice were examined for ocular structure under mild anesthesia with a ketamine-xylazine mixture using a slit lamp and Optical Coherence Tomography (OCT) (Envisu R-class, Leica). Following euthanasia and enucleation, eye size was measured with a caliper, and histological analysis was performed on 4 μm-thick hematoxylin and eosin (HE) stained sections from paraffin-embedded eyes fixed in Davidson’s solution. Ophthalmoscopic and histological ocular anomalies, including eye size shortening, cornea and lens abnormalities, retinal defects, and other abnormalities, were systematically recorded.

To assess the impact of different genotypes on the penetrance and severity of ocular anomalies, statistical analyses were conducted using R software (version 4.4.1). Logistic regression model using Firth’s bias reduction method (logistf package of R software) evaluated the association between genotype and phenotype, with “phenotype” as a binary variable (absent/present) and “genotype” as a factor with 6 levels (*Foxe3+/+*, *Foxe3*+/-, *Foxe3rv/+*, *Foxe3rv/rv*, *Foxe3rv/-, Foxe3-/-*). Odds ratios and 95% confidence intervals were calculated. Statistical significance was set at p < 0.05 and additional analyses were performed to evaluate the effects of age and gender on phenotype occurrence, and include them as covariates if necessary (**Table 1**).

To assess the association between the phenotype and both the number and nature of mutated alleles, a logistic regression adjusted for age (in months) was employed. The severity of the genotype was scored based on the number of mutated alleles and the severity of the mutation (allele + = 0, allele rv = 0.75 and allele - = 1; **Table 2)**, while phenotypic severity was classified into four levels for analysis: no phenotype (0), mild (cataract; 1), moderate (anterior segment dysgenesis; 2) and severe (significant developmental anomalies with eye growth defects; 3) (**Table 2**). An ordinal regression model using cumulative link models (CLMs, ordinal package in R software) was applied to analyze the severity of the phenotype in mice based on the number and severity of mutated alleles (**Table 3**).

### Assessing Ocular Abnormalities in Developing Mouse Embryos

For analysis of ocular development during embryonic life, mice were bred at 6 pm and checked for a vaginal plug 16 hours later. Females with a plug and a weight gain of more than 3g over 7 days were euthanized at embryonic (E) days 12.5, 13.5, 14.5 or 18.5, or at birth (P0) to collect embryos. Embryos were fixed in 10% buffered formalin at 4°C for 24 hours, heads were separated, embedded in paraffin and 4 µm thick eye tissue sections were subjected to HE staining. Ocular anomalies were systematically recorded.

### Quantitative Real-Time PCR analysis of Foxe3 expression in mouse lens epithelium

Mice aged 2 to 5 months were euthanized, and their eyes were enucleated and dissected on ice to isolate the lens epithelium. The procedure involved opening the eyes at the ora serrata, removing the lens, and carefully separating the capsule from the fiber mass. The lens epithelia from both eyes were pooled and total RNA was extracted using the RNeasy Mini Kit (Qiagen, Hilden, Germany), following the manufacturer’s protocol. RNA purity and concentration were assessed using a spectrophotometer (Nanodrop, Thermo Fisher Scientific, Waltham, MA, USA). mRNA was reverse transcribed from 35 ng of total RNA using the cDNA Verso Kit (Thermo Fisher Scientific) according to the manufacturer’s instructions. The abundance of *Foxe3* mRNA, along with the housekeeping genes *Tbp, Gusb, Gapdh,* and *Hprt1,* was quantified by real-time qPCR using specific primers **(Table S3)** and the SsoAdvanced Universal SYBR Green Supermix (Bio-Rad, Hercules, CA, USA) in a Realplex2 Mastercycler (Eppendorf, Montesson, France). Relative expression of *Foxe3* was calculated using the method described by Vandesompele et al. (Vandesompele et al. 2002) and compared across samples using a t-test or other appropriate statistical tests.

### Western blot analysis of FOXE3 in mouse lens epithelium (capsule)

Mouse lens epithelium were prepared as described above. Proteins (70 μg) were prepared using RIPA lysis buffer (ThermoFisher Scientific) and resolved on a Mini-ProteanTGX Stain Free 4-15% gel according to the supplier’s recommendations (Bio-Rad, Marne la Coquette, France). Proteins were transferred to a PVDF membrane (Biorad) using a RTA transfer kit (Bio-Rad) which was probed with FOXE3 (1:100; Santa Cruz Biotechnology, sc-377465) and Vinculin (1:2000; Abcam, ab91459). Secondary antibodies, either mouse IgG or rabbit IgG HRP (Thermo Fisher Scientific), were used at a 1:8000 dilution. Membranes were developed with Clarity Western ECL substrate (Bio-Rad) and visualized using a Chemidoc MP Imaging System (Bio-Rad). Protein quantification was performed using the gel analysis tool in Fiji (v. 1.53t) software.

### Analysis of the spatiotemporal expression patterns of Foxe3 and three candidate genes identified through pull-down assays

Co-expression of *Foxe3* with *Usf2*, *Cnbp* and *Gabpa* during ocular development was analyzed using RNAscope to visualize cell-specific expression and tissue morphology. E12.5 to E18.5 embryos were fixed in 4% PFA at 4°C for 24 hours, followed by washing in PBS. After embedding in OCT compound, the heads were rapidly frozen in liquid nitrogen and stored at - 80°C. For sectioning, heads were equilibrated to -20°C for 1 hour and sectioned at 14 µm thickness using a Cryostat set to -20°C for the object and -18°C for the chamber. Sections were air-dried for 1-2 hours and stored at -80°C.

Slides were prepared for RNAscope analysis according to the manufacturer’s instructions for frozen sections (Advanced Cell Diagnostics, Biotechne, Rennes, France). In brief, the sections were dehydrated in ethanol baths, dried, and treated with H₂O₂ for 10 minutes at room temperature. Proteinase digestion was performed the following day with Protease IV for 20 minutes at room temperature. Head sections were incubated for 2 hours at 40°C with a probe mix targeting *Foxe3* and the candidate genes *Usf2, Cnbp,* and *Gabpa*. Positive controls included probes for *Polr2a, Ppib*, and *Ubc* while *DapB* served as the negative control for the experimental groups. RNAscope signal amplification, including a series of amplifiers, HRP activators, blockers, and fluorophore incubations at 40°C, was conducted following the manufacturer’s protocol. Opals (Akoya Bioscience) were diluted 1:1500 in TSA buffer and assigned to channels C1 (Opal 570), C2 (Opal 690), and C3 (Opal 520). Nuclei were stained using DAPI, and the sections were mounted with Fluoromount, and stored at -20°C for at least 12 hours before imaging.

Images were acquired with a Zeiss Spinning Disk Confocal. Whole sections were scanned using a 10X objective, and detailed images of lens epithelium were captured with a 40X or 63X oil objective as maximal projections of 10 or 30 stacks covering 6.09 µm (frozen sections). Images were saved as .czi files and converted to TIFF format for analysis. Quantitative image analysis was performed using Fiji (v. 1.53t) and CellProfiler (v. 4.2.1) software following the analysis method suggested by the manufacturer. In brief, the presumptive lens was manually defined as the ROI. The integrated intensity of the ROI was measured for each channel, along with the area covered by the DAPI-signal. For each channel, 25 spots were measured to determine the average intensity value of one mRNA spot, and 25 nuclei were delimited to measure the average area of one nucleus. Dividing the integrated intensity of the ROI by the average dot intensity estimates the total dots of mRNA in the region, and this value can be reported to the number of nuclei, resulting in a mean value of dots/nuclei. Unpaired bilateral Student’s t-test was used for two-group analyses. Data are presented as the mean ± SEM; p<0.05 was considered significant.

### siRNA-mediated knockdown of Usf2, Cnbp and Gabpa expression in cultured epithelial lens cells from Foxe3 mouse lines

*Foxe3 +/+* and *rv/rv* mouse between 2 and 4 months of age were euthanized, and their eyes were enucleated. Eyes were briefly washed with 70% ethanol and incubated in PBS under sterile conditions. Lens capsule were recovered as described above and anterior epithelial cells were incubated in P12 cell culture plates with EpiCM at 37°C with 5% CO₂ until the first cells adhered to the plate, typically within 48 hours. Once the cells reached confluence, they were maintained in EpiCM on plates coated with poly-L-lysine (Sigma-Aldrich; Saint-Quentin-Fallavier, France) and passaged using 0.05% trypsin-EDTA (Thermo Fisher Scientific).

For siRNA knock-down, cells at 70% confluence were transfected with *Usf2, Cnbp and Gabpa* (50 nM; FlexiTube GeneSolution,Qiagen; Courtaboeuf, France, Gene ID: GS22282, GS12787 and GS14390 respectively) or a no-target siRNAs (AllStars Neg. Control siRNA, Qiagen; Cat#1027281) using Lipofectamine 2000 reagent (Invitrogen, Illkirch, France). Transfected cells were harvested after 48 hours and total RNA was extracted for real-time qPCR analysis as described above using specific primers (**Table S3).** *Foxe3* data represent the mean ± SD of four independent experiments, normalized to the no-target control siRNA, with statistical significance evaluated using a two-way ANOVA followed by Dunnett’s post-hoc test.

## DATA AND CODE AVAILABILITY

The mass spectrometry proteomics data have been deposited to the ProteomeXchange Consortium via the PRIDE (Perez-Riverol et al. 2022) partner repository with the dataset identifier PXD058162.

## DECLARATION OF INTEREST

None

## ACKNOWLEDGEMENTS

This research has been generously supported by grants from the “Institut National de la Santé et de la Recherche Médicale” (INSERM), MSDAVENIR (DEVO-DECODE program), the “Fondation Visio”, and the “Association Retina France”.

## AUTHOR CONTRIBUTIONS

JP, NC, JMR and LFT designed the project. PD generated the MCOR mice. JP, LV, CA and EE performed and interpreted the molecular, histological and imaging experiments. DH analyzed the Hi-Cseq data. JP, CA, IRL, MG, J-YD and YAM, performed ophthalmological and histological studies in mice. LV performed *in vivo* experiments. CV-D, PC, JP and NCh provided clinical data. IK, MV and MB performed DNA pull-down experiments. FJ-H and FL performed biostatistics analyses. JP wrote the manuscript and prepared the figures with contributions from all coauthors. NC, JMR and LFT reviewed all of the data and edited the manuscript.

## DECLARATION OF GENERATIVE AI AND AI-ASSISTED TECHNOLOGIES IN THE WRITING PROCESS

During the preparation of this work the author(s) used ChatGPT in order to edit English. After using this tool/service, the authors reviewed and edited the content as needed and take full responsibility for the content of the publication.

## Notes

### Competing Interest Statement

The authors have declared no competing interest.

