## Supplemental data for "Insights into the FOXE3 Transcriptional Network and Disease Mechanisms from the Investigation of a Regulatory Variant Driving Complex Microphthalmia"

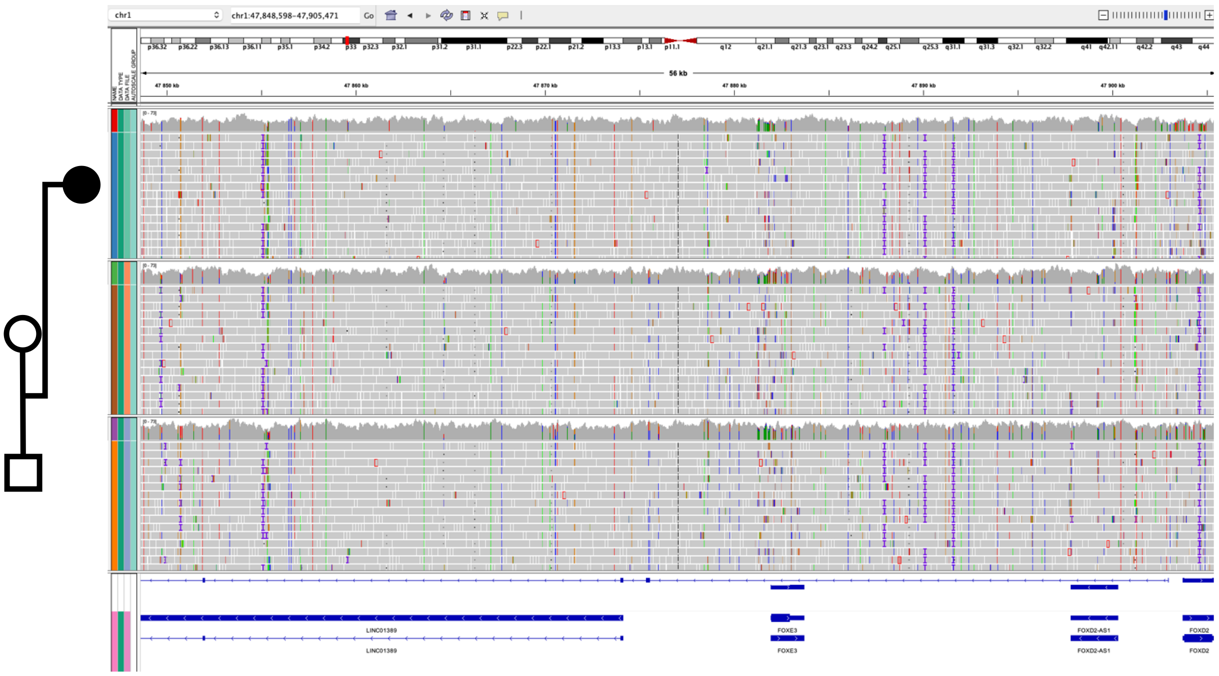


**Figure S1. Whole-genome sequencing read alignments of the affected individual and her parents.** Visualization of read alignments using the Integrated Genomics Viewer (IGV) demonstrates the absence of structural variant in the genomic region surrounding *FOXE3*.

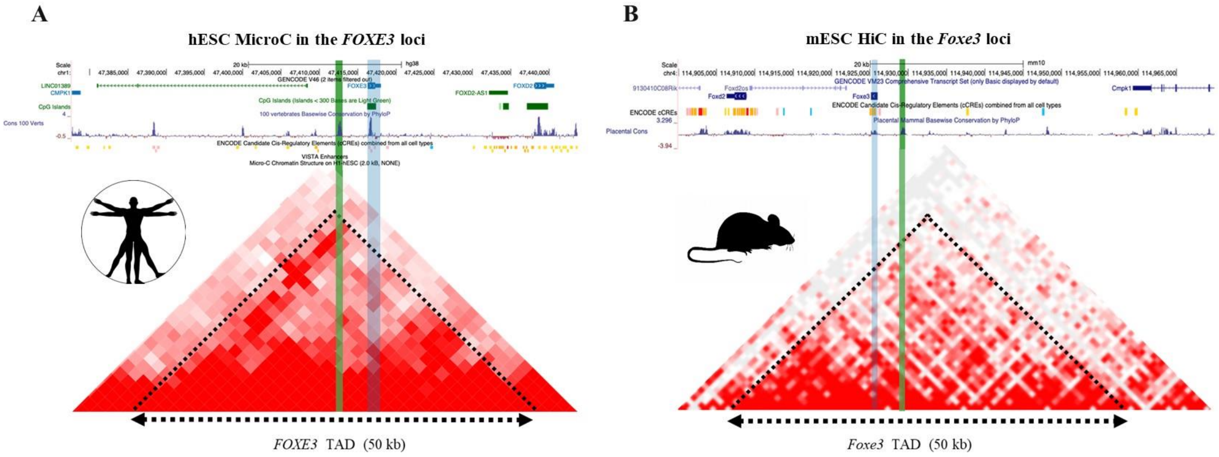


**Figure S2: Structural organization of the *FOXE3* region in human and mouse embryonic stem cells.** (A) MicroC of human chromosome 1p33 (GRCh38/hg38 chr1:47,385,000-47,435,000 and (B) Hi-C map of mouse chromosome 4qD1(GRCm38/mm10_chr4: 114,910,000-114,960,000) encompassing the *FOXE3* locus demonstrate that the locus resides within a 50 kb topologically associating domain (TAD) in both human and mouse embryonic stem cells (hESCs and mESCs, respectively). The positions of *FOXE3* and the regulatory variant are marked in blue and green, respectively. Notably, the entire region is inverted in the mouse genome compared to the human genome.

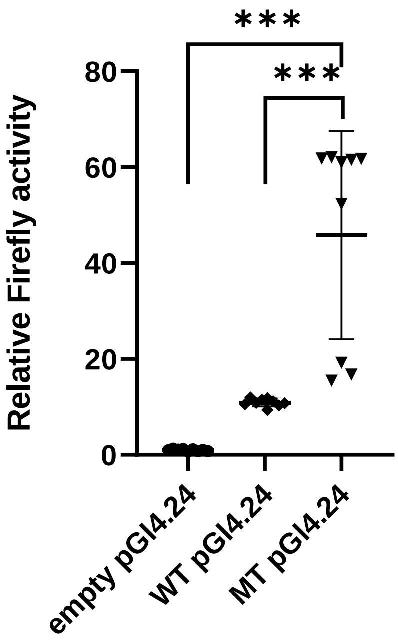


**Figure S3. *In vitro* luciferase assay analysis of wild-type and mutant *FOXE3* sequences.** Luciferase assays demonstrate 10-fold and 45-fold increase in firefly luciferase activity for the wild-type (WT) and mutant (MT) *FOXE3* sequences expressed in the pGL4.24 vector, respectively, compared to the empty vector. *** = $p < 0.0001$

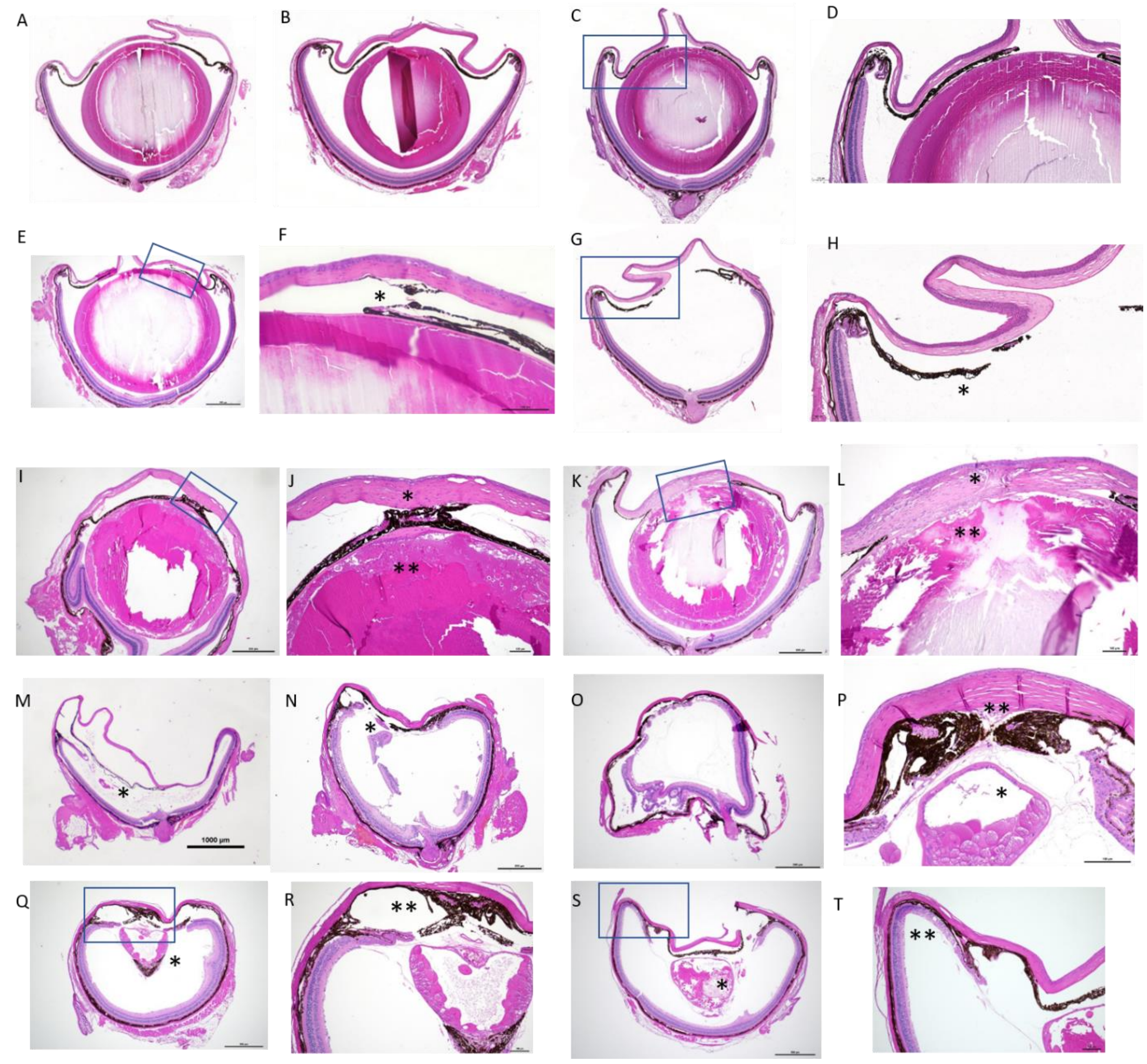


**Figure S4. Histological analysis of the eyes from adult *Foxe3* wild-type and mutant animals.**
(A-B) Histological images of the left (A) and right (B) eyes from a *Foxe3+/+* animal (Specimen 430; **Table S4**), showing the typical spherical lens that occupies a large portion of the posterior segment, as described by Chalupa (2008).
(C-D) Histological image from a *Foxe3+/-* animal (specimen 379; **Table S4**), showing normal ocular structure (C, magnified in D).
(E-H) Histological images from two *Foxe3rv/+* animals (452 and 453 specimens; **Table S4**) showing extensive focal irido-corneal adhesion (*) (E, magnified in F) and normal ocular structure (G, magnified in H; the lens is not visible due to a technical artifact, indicated by nonspecific vacuolization (*) of the iris posterior epithelium), respectively.
(I-L) Histological images from two *Foxe3rv/-* mice (328 and 373 specimens; **Table S4**) showing cataractic lenses (**), extensive focal irido-lenticular and irido-corneal adhesions (*), and focal fibrosis with thickening of the anterior lens capsule at the adhesion site in the right eye (I, magnified in J), and chronic keratitis with a central transcorneal lesion (*), focal fibrosis, and adherence between the cornea and the anterior capsule of the cataractic lens (**) (K, magnified in L), respectively.

**M-T.** Histological analysis of the left and right eyes from three *Foxe3-/-* animals (315, 330 and 349 specimens; **Table S4**) showing irido-corneal adhesion, posterior iris cystic formation, lens remnants (*), and proteic strands and flakes surrounding the residual lens and within the vitreous in the left eye (M), irido-corneal adhesion and lens remnants (*) in the right eye (N), irido-ciliary and corneal adhesions, absence of the lens, retinal detachment, rosette-like retinal structures, and tombstone-like hypertrophy of the retinal pigment epithelium in the right eye (O), extensive fibrous adhesions between the cornea, iris and lens (**) with a small, irregularly shaped lens (*) with a prominent anterior empty vesicle and cataractic swollen fibers (P), extensive uveo-corneal adhesions (**) and a triangular, small cataractic lens (*) with large anterior vesiculation, anteriorly displaced retina, retinal detachment, and retinal folds in the right eye (Q, magnified in R) and uveo-corneal adhesions (**) and a vacuolated triangular cataractic lens (*) in the left eye (S, magnified in T), respectively.

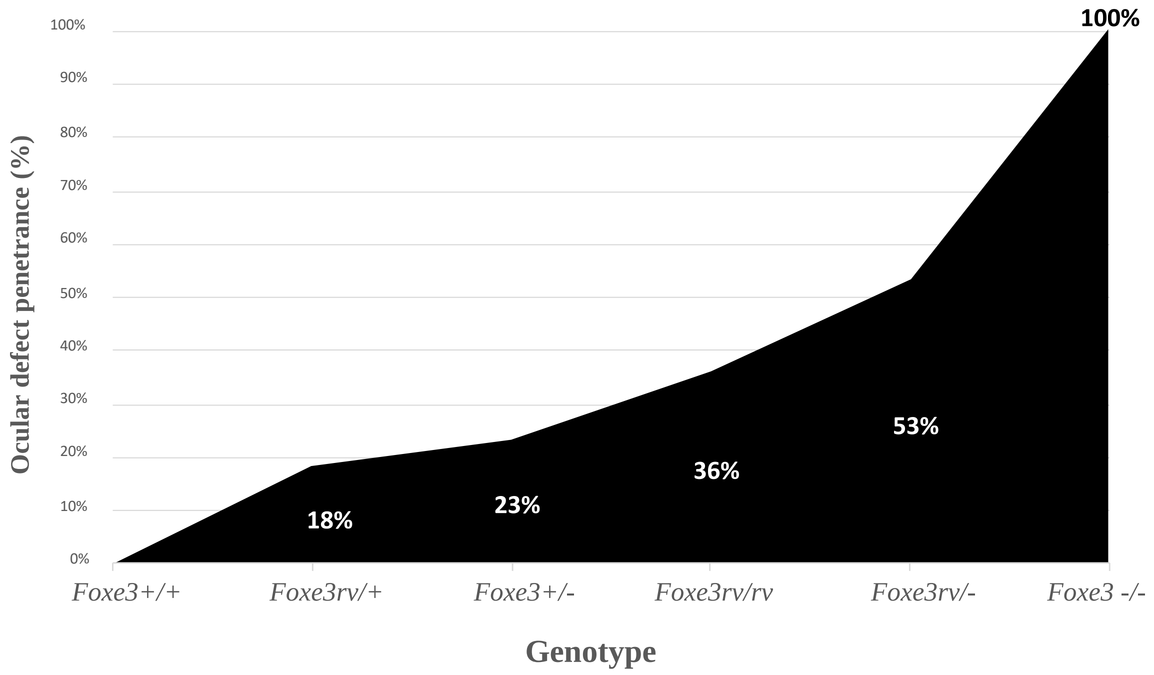


**Figure S5. Ocular defect penetrance by genotype.** The increasing trend in ocular defect penetrance correlates with the escalating severity of *Foxe3* genotypes. Data points are annotated with percentages, representing the exact penetrance observed for each genotype.

**Table S1. Oligonucleotides designed for the wild-type and mutant human non-coding sequences were used in DNA pull-down and *in vitro* luciferase assays.** The wildtype and mutant rs745674596 nucleotides are highlighted in bold and enlarged text. BglI cohesive ends used for cloning are underlined.

| | Sequence (5'-3') |
|---|---|
| *FOXE3*-WT_F | TTCACTGGCCATCTTATGCCCGGATGCTCAGCCGCATCAC**G**TCCGGCCCAGG<br>GCCTGTGAAAAGAGGGCCCAGCCACGCTGCGG |
| *FOXE3*-WT _R | CAGCGTGGCTGGGCCCTCTTTTCACAGGCCCTGGGCCGGA**C**GTGATGCGGCT<br>GAGCATCCGGGCATAAGATGGCCAGTGAAGTT |
| *FOXE3*-Mut_F | TTCACTGGCCATCTTATGCCCGGATGCTCAGCCGCATCAC**A**TCCGGCCCAGG<br>GCCTGTGAAAAGAGGGCCCAGCCACGCTGCGG |
| *FOXE3*-Mut_R | CAGCGTGGCTGGGCCCTCTTTTCACAGGCCCTGGGCCGGA**T**GTGATGCGGCT<br>GAGCATCCGGGCATAAGATGGCCAGTGAAGTT |

**Table S2. Primers used to verify the introduction of the variants and for genotyping the *Foxe3 rv* and *Foxe3-* alleles in the mouse.**

| | Forward (5'-3') | Reverse (5'-3') |
|---|---|---|
| *Foxe3 rv* | TATCCACCCAGTGTGGTCAG | AGGCCATGTGCATAGTCTGG |
| *Foxe3-* | CACCACTTCCTCCGGTTCG | ATATAAGGCGAACCTGCGACC |

**Table S3. A comprehensive list of oligonucleotides for RT-qPCR analysis.**

| | Forward (5'-3') | Reverse (5'-3') |
|---|---|---|
| *Foxe3* | GCCGCCCTACTCATACATC | ACAGTCGTTGAGGGTGAGG |
| *Tbp* | TGACCTAAAGACCATTGCACTTCGT | CTGCAGCAAATCGCTTGGGA |
| *Hprt1* | GTTGGTACAGGCCAGACTTTGTT | AAACGTGATTCAAATCCCTGAAGTA |
| *Gusb* | CTGGGGTTGTGATGTGGTCTGT | TGTGGGTGATCAGCGTCTTAAAGT |
| *Gapdh* | TTCACCACCATGGAGAAGGC | GGCATGGACTGTGGTCATGA |
| *Cnbp* | TCTTCGTCCGAGTCTCCTCC | GGAAACAAACTGGAACCCTCT |
| *Usf2* | CAATGAAGTGGAACGGAGAAG | TCCCGGATGTAATCGCAAG |
| *Gabpa* | CCGGGGAACAGAACAGGAAA | TTTTGCGCAACCAACTCAGG |

**Table S4. Phenotypic analysis of *Foxe3+/+, Foxe3+/rv, Foxe3+/-, Foxe3rv/rv, Foxe3rv/-,* and *Foxe3-/-* genotypes.** The different genotypes were assessed and compared for ocular globe size and structural integrity using caliper measurements and slit lamp examination. **SN:** Specimen number, **F:** female**, M:** male**, Age:** age at examination, **LE: l**eft eye, **RE:** right eye.

| SN | Sex | Age (months) | Genotype | LE (size in mm) | RE (size in mm) | Ophthalmological phenotype (slit lamp) |
|---|---|---|---|---|---|---|
| 376 | F | 8 | *Foxe3 rv/+* | 3,5 | 4 | Bilateral central subcapsular cataract |
| 384 | M | 8 | *Foxe3 rv/+* | 4 | 4 | Normal |
| 385 | F | 8 | *Foxe3 rv/+* | 3,5 | 3,5 | Normal |
| 386 | F | 8 | *Foxe3 rv/+* | 3,5 | 3,5 | Normal |
| 387 | F | 8 | *Foxe3 rv/+* | 3,5 | 3,5 | Normal |
| 402 | F | 7 | *Foxe3 rv/+* | 3,5 | 4 | Bilateral nuclear cataract |
| 404 | F | 7 | *Foxe3 rv/+* | 4 | 3,5 | Normal |
| 405 | F | 7 | *Foxe3 rv/+* | 3,5 | 3,5 | Normal |
| 431 | M | 6 | *Foxe3 rv/+* | 3,5 | 3,5 | Normal |
| 433 | M | 6 | *Foxe3 rv/+* | 4 | 4 | Normal |
| 435 | M | 6 | *Foxe3 rv/+* | 3,5 | 3,5 | Normal |
| 436 | M | 6 | *Foxe3 rv/+* | 4 | 4 | Normal |
| 437 | F | 6 | *Foxe3 rv/+* | 3,5 | 4 | Normal |
| 445 | M | 5 | *Foxe3 rv/+* | 3,5 | 3,5 | Normal |
| 447 | M | 5 | *Foxe3 rv/+* | 4 | 4 | Normal |
| 449 | M | 5 | *Foxe3 rv/+* | 4 | 4 | Normal |
| 450 | M | 5 | *Foxe3 rv/+* | 3,5 | 3,5 | Dyscoria, irido-corneal adherence with corneal clouding (LE) |
| 451 | M | 5 | *Foxe3 rv/+* | 4 | 4 | Normal |
| 452 | M | 5 | *Foxe3 rv/+* | 3,5 | 3,5 | Irido-corneal adherence with corneal clouding (RE) |
| 453 | F | 5 | *Foxe3 rv/+* | 3,5 | 3,5 | Normal |
| 454 | F | 5 | *Foxe3 rv/+* | 4 | 3,5 | Normal |
| 464 | F | 4 | *Foxe3 rv/+* | 3,5 | 3,5 | Normal |
| 465 | F | 4 | *Foxe3 rv/+* | 4 | 4 | Normal |
| 466 | F | 4 | *Foxe3 rv/+* | 3,5 | 3,5 | Normal |
| 467 | F | 4 | *Foxe3 rv/+* | 3,5 | 3,5 | Small irido-corneal adherence (RE) |
| 468 | M | 4 | *Foxe3 rv/+* | 3,5 | 3,5 | Normal |
| 469 | M | 4 | *Foxe3 rv/+* | 4 | 4 | Normal |
| 482 | F | 4 | *Foxe3 rv/+* | 3,5 | 3,5 | Normal |
| 309 | M | 11 | *Foxe3 rv/rv* | 4 | 4 | Normal |
| 310 | M | 11 | *Foxe3 rv/rv* | 4 | 4 | Normal |
| 313 | F | 11 | *Foxe3 rv/rv* | 3,5 | 3,5 | Normal |
| 325 | M | 11 | *Foxe3 rv/rv* | 4 | 4 | Normal |
| 327 | M | 11 | *Foxe3 rv/rv* | 3,5 | 4 | Normal |
| 331 | F | 11 | *Foxe3 rv/rv* | 4 | 4 | Anterior subcapsular cataract (LE) |
| 336 | M | 10 | *Foxe3 rv/rv* | 4 | 3,5 | Anterior subcapsular cataract (LE) |
| 338 | M | 10 | *Foxe3 rv/rv* | 4 | 4 | Normal |
| 347 | F | 10 | *Foxe3 rv/rv* | 4 | 4 | Normal |
| 350 | M | 8 | *Foxe3 rv/rv* | 3,5 | 4 | Anterior subcapsular cataract (LE) |
| 352 | M | 8 | *Foxe3 rv/rv* | 4 | 4 | Nuclear cataract (LE) |
| 354 | M | 8 | *Foxe3 rv/rv* | 4 | 3,5 | Normal |
| 355 | F | 8 | *Foxe3 rv/rv* | 3,5 | 3,5 | Nuclear cataract (LE) |
| 391 | F | 8 | *Foxe3 rv/rv* | 3,5 | 3,5 | Normal |
| 392 | M | 7 | *Foxe3 rv/rv* | 3,5 | 3,5 | Normal |
| 397 | M | 7 | *Foxe3 rv/rv* | 4 | 4 | Normal |
| 407 | M | 7 | *Foxe3 rv/rv* | 4 | 4 | Normal |
| 420 | M | 7 | *Foxe3 rv/rv* | 4 | 4 | Normal |
| 428 | M | 6 | *Foxe3 rv/rv* | 4 | 4 | Normal |
| 478 | M | 4 | *Foxe3 rv/rv* | 3,5 | 3,5 | Normal |

| | | | | | | |
|---|---|---|---|---|---|---|
| 229 | M | 11 | *Foxe3 rv/rv* | 3,5 | 3,5 | Normal |
| 230 | F | 11 | *Foxe3 rv/rv* | 3,5 | 3,5 | Normal |
| 232 | F | 11 | *Foxe3 rv/rv* | 3,5 | 3,5 | Irido-corneal adherence with corneal clouding and cataract (RE) |
| 242 | F | 8 | *Foxe3 rv/rv* | 3,5 | 3,5 | Bilateral cataract |
| 243 | F | 8 | *Foxe3 rv/rv* | 3,5 | 3,5 | Cataract (LE) |
| 251 | F | 7 | *Foxe3 rv/rv* | 3,5 | 3,5 | Normal |
| 253 | F | 7 | *Foxe3 rv/rv* | 3,5 | 3,5 | Cataract (RE) |
| 255 | F | 7 | *Foxe3 rv/rv* | 4 | 3,5 | Bilateral cataract |
| 326 | M | 11 | *Foxe3 rv/-* | 3,5 | 3,5 | Normal |
| 328 | F | 11 | *Foxe3 rv/-* | 4 | 3,5 | Iris coloboma (LE); corneal clouding (1 central et 1 peripheral) with reduced anterior chamber and total cataract (RE) |
| 329 | F | 11 | *Foxe3 rv/-* | 4 | 4 | Bilateral cataract |
| 348 | F | 10 | *Foxe3 rv/-* | 4 | 4 | Cataract (LE) |
| 364 | F | 8 | *Foxe3 rv/-* | 3,5 | 4 | Normal |
| 371 | F | 8 | *Foxe3 rv/-* | 4 | 4 | Normal |
| 372 | F | 8 | *Foxe3 rv/-* | 3,5 | 4 | Normal |
| 373 | F | 8 | *Foxe3 rv/-* | 3,5 | 3,5 | Central pit with retrocorneal lens, reduced anterior chamber (RE) |
| 393 | M | 7 | *Foxe3 rv/-* | 4 | 4 | Dyscoria with irido-corneal and irido-lenticular adherences (LE) |
| 411 | M | 7 | *Foxe3 rv/-* | 3,5 | 4 | Bilateral anterior capsular cataract |
| 412 | M | 7 | *Foxe3 rv/-* | 4 | 3,5 | Normal |
| 416 | M | 7 | *Foxe3 rv/-* | 4 | 4 | Normal |
| 417 | M | 7 | *Foxe3 rv/-* | 3,5 | 3,5 | Anterior capsular cataract (RE) |
| 418 | M | 7 | *Foxe3 rv/-* | 4 | 4 | Mild dyscoria (LE) |
| 423 | F | 7 | *Foxe3 rv/-* | 3,5 | 3,5 | Bilateral polar posterior cataract |
| 456 | F | 4 | *Foxe3 rv/-* | 4 | 4 | Normal |
| 457 | F | 4 | *Foxe3 rv/-* | 3,5 | 3,5 | Normal |
| 461 | M | 4 | *Foxe3 rv/-* | 3,5 | 3,5 | Iris coloboma (LE) |
| 477 | M | 4 | *Foxe3 rv/-* | 4 | 4 | Normal |
| 479 | M | 4 | *Foxe3 rv/-* | 3,5 | 3,5 | Normal |
| 480 | M | 4 | *Foxe3 rv/-* | 3,5 | 3,5 | Normal |
| 481 | M | 4 | *Foxe3 rv/-* | 3,5 | 3,5 | Normal |
| 483 | F | 4 | *Foxe3 rv/-* | 3,5 | 3,5 | Normal |
| 244 | F | 8 | *Foxe3 rv/-* | | | Bilateral cataract |
| 264 | F | 5 | *Foxe3 rv/-* | 3,5 | 3,5 | Normal |
| 231 | F | 11 | *Foxe3 rv/-* | 3,5 | 3,5 | Corneal clouding, small lens, reduced anterior chamber |
| 233 | M | 9 | *Foxe3 rv/-* | 3,5 | 3,5 | Bilateral cataract |
| 234 | M | 9 | *Foxe3 rv/-* | 3,5 | 3,5 | Bilateral cataract |
| 235 | M | 9 | *Foxe3 rv/-* | 3,5 | 3,5 | Normal |
| 237 | F | 8 | *Foxe3 rv/-* | 3,5 | 3,5 | Cataract (LE) |
| 239 | M | 8 | *Foxe3 rv/-* | 3,5 | 3,5 | Normal |
| 246 | F | 8 | *Foxe3 rv/-* | 3,5 | 3,5 | Nuclear cataract (RE) |
| 256 | M | 5 | *Foxe3 rv/-* | 3,5 | 3,5 | Normal |
| 258 | M | 5 | *Foxe3 rv/-* | 3,5 | 3,5 | Normal |
| 260 | F | 5 | *Foxe3 rv/-* | 3,5 | 3,5 | Cataract (LE) |
| 261 | F | 5 | *Foxe3 rv/-* | 3,5 | 3,5 | Normal |
| 262 | M | 5 | *Foxe3 rv/-* | 3,5 | 3,5 | Bilateral cataract |
| 263 | M | 5 | *Foxe3 rv/-* | 4 | 4 | Cataract (RE) |
| 266 | M | 5 | *Foxe3 rv/-* | 4 | 4 | Bilateral cataract |
| 267 | F | 5 | *Foxe3 rv/-* | 3,5 | 3,5 | Coloboma (RE) |
| 377 | F | 8 | *Foxe3 +/-* | 3,5 | 4 | Cataract (RE) |
| 378 | F | 8 | *Foxe3 +/-* | 3,5 | 3,5 | Normal |
| 379 | F | 8 | *Foxe3 +/-* | 3,5 | 3,5 | Normal |
| 380 | M | 8 | *Foxe3 +/-* | 4 | 4 | Normal |

| | | | | | | |
|---|---|---|---|---|---|---|
| 381 | M | 8 | *Foxe3* +/- | 3,5 | 3,5 | Normal |
| 382 | M | 8 | *Foxe3* +/- | 4 | 4 | Normal |
| 383 | M | 8 | *Foxe3* +/- | 4 | 4 | Bilateral dyscoria |
| 401 | M | 7 | *Foxe3* +/- | 4 | 4 | Normal |
| 403 | F | 7 | *Foxe3* +/- | 3,5 | 3,5 | Normal |
| 406 | F | 7 | *Foxe3* +/- | 3,5 | 3,5 | Normal |
| 432 | M | 6 | *Foxe3* +/- | 3,5 | 3,5 | Coloboma (RE) |
| 434 | M | 6 | *Foxe3* +/- | 3,5 | 3,5 | Normal |
| 438 | F | 6 | *Foxe3* +/- | 4 | 4 | Normal |
| 439 | F | 6 | *Foxe3* +/- | 3,5 | 3,5 | Iridolenticular adherence (LE) |
| 444 | M | 5 | *Foxe3* +/- | 3,5 | 3,5 | Anterior capsular cataract |
| 446 | M | 5 | *Foxe3* +/- | 3,5 | 4 | Normal |
| 448 | M | 5 | *Foxe3* +/- | 4 | 4 | Normal |
| 459 | M | 4 | *Foxe3* +/- | 4 | 4 | Normal |
| 460 | M | 4 | *Foxe3* +/- | 4 | 4 | Normal |
| 462 | F | 4 | *Foxe3* +/- | 3,5 | 4 | Normal |
| 463 | F | 4 | *Foxe3* +/- | 3,5 | 3,5 | Normal |
| 475 | F | 4 | *Foxe3* +/- | 4 | 4 | Normal |
| 315 | M | 11 | *Foxe3* -/- | 3 | 2,5 | Bilateral microphthalmia with total corneal clouding, athalamia more severe (RE) |
| 316 | F | 11 | *Foxe3* -/- | 3 | 2,5 | Bilateral microphthalmia with total corneal clouding, athalamia, total adherence of the iris against the cornea on both eyes |
| 317 | F | 11 | *Foxe3* -/- | 1,5 | 1,5 | Severe bilateral microphthalmia |
| 330 | F | 11 | *Foxe3* -/- | 3 | 3 | Bilateral microphthalmia with total corneal clouding, athalamia |
| 349 | F | 10 | *Foxe3* -/- | 3 | 2,5 | Bilateral microphthalmia with total corneal clouding, central pit, athalamia |
| 419 | M | 7 | *Foxe3* -/- | 2 | 2 | Bilateral microphthalmia with total corneal clouding (RE), athalamia (LE) |
| 422 | F | 7 | *Foxe3* -/- | 2 | 2,5 | Bilateral microphthalmia |
| 455 | M | 4 | *Foxe3* -/- | 2,5 | 2 | Bilateral microphthalmia with total corneal clouding, athalamia |
| 458 | F | 4 | *Foxe3* -/- | 2,5 | 1,5 | Bilateral microphthalmia with total corneal clouding, athalamia |
| 476 | M | 4 | *Foxe3* -/- | 2,5 | 2,5 | Bilateral microphthalmia with total corneal clouding, athalamia, central pit (LE) |
| 236 | F | 9 | *Foxe3* -/- | 3 | 3 | Bilateral microphthalmia with cataract (LE) and corneal clouding, athalamia, ectopia lentis (RE) |
| 265 | M | 5 | *Foxe3* -/- | 3 | 3 | Bilateral microphthalmia with total corneal clouding, athalamia, ectopia lentis on both eye |
| 408 | F | 8 | *Foxe3* +/+ | 3,5 | 3,5 | Normal |
| 409 | F | 8 | *Foxe3* +/+ | 3,5 | 3,5 | Normal |
| 410 | F | 8 | *Foxe3* +/+ | 3,5 | 3,5 | Normal |
| 411 | F | 8 | *Foxe3* +/+ | 4 | 4 | Normal |
| 420 | F | 7 | *Foxe3* +/+ | 3,5 | 3,5 | Normal |
| 421 | F | 7 | *Foxe3* +/+ | 4 | 4 | Normal |
| 427 | F | 6 | *Foxe3* +/+ | 3,5 | 3,5 | Normal |
| 428 | F | 6 | *Foxe3* +/+ | 3,5 | 3,5 | Normal |
| 429 | F | 6 | *Foxe3* +/+ | 4 | 4 | Normal |
| 430 | F | 6 | *Foxe3* +/+ | 4 | 4 | Normal |
| 450 | F | 5 | *Foxe3* +/+ | 3,5 | 3,5 | Normal |
| 451 | F | 5 | *Foxe3* +/+ | 3,5 | 3,5 | Normal |
| 608 | M | 7 | *Foxe3* +/+ | 4 | 4 | Normal |
| 610 | M | 7 | *Foxe3* +/+ | 4 | 4 | Normal |
| 633 | M | 5 | *Foxe3* +/+ | 3,5 | 4 | Normal |
| 634 | M | 5 | *Foxe3* +/+ | 3,5 | 4 | Normal |

| | | | | | | |
|---|---|---|---|---|---|---|
| 102 | M | 6 | *Foxe3* +/+ | 3,5 | 3,5 | Normal |
| 110 | M | 6 | *Foxe3* +/+ | 4 | 4 | Normal |
| 134 | M | 4 | *Foxe3* +/+ | 3,5 | 3,5 | Normal |
| 135 | M | 4 | *Foxe3* +/+ | 3,5 | 3,5 | Normal |
| 137 | M | 4 | *Foxe3* +/+ | 3,5 | 3,5 | Normal |
| 143 | F | 4 | *Foxe3* +/+ | 3,5 | 3,5 | Normal |
| 144 | F | 4 | *Foxe3* +/+ | 3,5 | 3,5 | Normal |
| 146 | F | 4 | *Foxe3* +/+ | 3,5 | 4 | Normal |
| 553 | M | 5 | *Foxe3* +/+ | 3,5 | 3,5 | Normal |
| 554 | M | 5 | *Foxe3* +/+ | 3,5 | 3,5 | Normal |
| 358 | M | 10 | *Foxe3* +/+ | 3,5 | 3,5 | Normal |
| 359 | M | 10 | *Foxe3* +/+ | 3,5 | 3,5 | Normal |
| 373 | F | 8 | *Foxe3* +/+ | 3,5 | 3,5 | Normal |
| 376 | F | 8 | *Foxe3* +/+ | 3,5 | 3,5 | Normal |
